# Programmable RNA base editing with a single gRNA-free enzyme

**DOI:** 10.1101/2021.08.31.458316

**Authors:** Wenjian Han, Wendi Huang, Miaowei Mao, Tong Wei, Yanwen Ye, Zefeng Wang

**Affiliations:** Bio-Med Big Data Center, CAS Key Laboratory of Computational Biology, CAS Center for Excellence in Molecular Cell Science, Shanghai Institute of Nutrition and Health, University of Chinese Academy of Sciences, Chinese Academy of Sciences, Shanghai 200031, China; University of Chinese Academy of Sciences, Beijing 100049, China

## Abstract

Programmable RNA editing enables rewriting gene expression without changing genome sequences. Current tools for specific RNA editing dependent on the assembly of guide RNA into an RNA/protein complex, causing delivery barrier and low editing efficiency. We report a new gRNA-free system, RNA editing with individual RNA-binding enzyme (REWIRE), to perform precise base editing with a single engineered protein. This artificial enzyme contains a human-originated programmable PUF domain to specifically recognize RNAs and different deaminase domains to achieve efficient A-to-I or C-to-U editing, which achieved 60-80% editing rate in human cells, with a few non-specific editing sites in the targeted region and a low level off-target effect globally. The RNA-binding domain in REWIREs was further optimized to improve editing efficiency and minimize off-target effects. We applied the REWIREs to correct disease-associated mutations and achieve both types of base editing in mice. As a single-component system originated from human proteins, REWIRE presents a precise and efficient RNA editing platform with broad applicability.

## INTRODUCTION

Base editing is a new genetic modification strategy that directly installs point mutations into the DNAs or RNAs without making pernicious breaks in nucleic acid chain (1–4). Directed base editing has provided new potentials in treating genetic diseases caused by point mutations, such as Duchenne muscular dystrophy (5) or Hutchinson–Gilford progeria syndrome (6). Compared to DNA manipulation, editing on RNAs is non-permanent and reversible, making it relative safer for *in vivo* application as a potential therapy (7–10). Previously several groups had used guided antisense RNA oligonucleotide to recruit adenosine deaminases from the ADAR (<u>a</u>denosine <u>d</u>eaminases <u>a</u>cting on <u>R</u>NA) family to transform the RNA base from adenosine to inosine (recognized as guanosine) (3,11–16). However, the formation of dsRNA region and the recruitment of ADARs to the target RNAs are hard to control. Recently, Zhang’s group developed two CRISPR-based systems using RNA-targeting Cas13 protein and ADAR2 to achieve base editing of the specific RNAs: the REPAIR used the active domain of ADAR2 to achieve programmable adenosine to inosine (A-to-I) editing, and the RESCUE employed a modified ADAR2 to enable additional cytidine to uridine (C-to-U) editing (4, 17). More recently, the Chi’s group used similar design to engineer two CRISPR-based editing systems, REPAIRx and CURE, that improve the target recognition with dCasRx (18, 19). To overcome the difficulty for *in vivo* delivery of a large CRISPR-Cas system, a comparable but more concise system, CRISPR-Cas-inspired RNA targeting system (CIRTS), has also been developed by Dickinson’s group to execute A-to-I editing on specific RNA targets (20).

All these engineered RNA base editing systems require the assembly of guide RNA (gRNA) and protein effector to form an RNA-protein complex for recognizing target RNAs. However, the cellular RNAs have much higher copy numbers than the genomic DNAs, the assembly and co-folding of the RNA-protein complex can be the rate-limiting step that reduce editing efficiency and hinder its application (21). Pairing of the gRNA and target RNA also leads to a relatively large editing window with potential “bystander” editing sites (4,17,18). Finally, because Cas proteins are encoded by bacterial genome, the pre-existing immunity in a large fraction of human population presents a looming concern for future applications of CRISPR-based systems (22, 23). Alternatively, cellular RNAs can be directly manipulated using engineered proteins that contain a programmable RNA-binding module and different functional modules (24). For example, the RNA binding scaffold of Pumilio and FBF homology (PUF) proteins contains eight repeat motifs, each can be reprogrammed to specifically bind any RNA base through interaction with the Watson–Crick edge (25–27). The PUF-based systems with different functional domains have been developed by several groups to detect cellular RNAs (28) or manipulate RNA splicing (29), translation (30), degradation (31) or methylation/demethylation (32).

Here we engineered a single component RNA editing tool, <u>R</u>NA <u>e</u>diting <u>w</u>ith individual <u>R</u>NA-binding <u>e</u>nzyme (REWIRE), which contains a programmable PUF domain (25, 33) to specifically bind RNAs and a functional domain from human ADARs to edit adenosine or APOBEC3A to edit cytidine (2, 34) (Figure S1A and S1B). Since the PUF domain can be reprogrammed to recognize almost any short 8-nucleotide (8-nt) RNA sequences (35, 36), the REWIRE system can theoretically be designed to install two key transitions (A-to-I or C-to-U) in any RNA. We further redesigned the PUF domain in REWIREs to enhance the specificity and the editing efficiency. As a simple human-originated protein without any RNA components, this system provides an easier and more practical system for basic research and gene therapy.

## MATERIAL AND METHODS

### Plasmid construction

DNA fragments encoding the deaminase domains of the human ADAR1 and ADAR2 were amplified from the cDNA of HEK 293T cells. The hyperactive variants of hADAR1/2 were generated with primers containing corresponding mutations (37, 38). All hADAR1/2 fragments were cloned into the pCI-Neo vector (Promega) to generate the pCI-ADAR vectors. Different versions of codon optimized PUF repeats from human PUM1 that recognize all four RNA bases were chemically synthesized (GENEWIZ). The eight or ten PUF repeats were assembled by PCR amplification and inserted to N-terminal of ADARs to generate various AI-REWIRE expression vectors. The same strategy was applied to generate the CU-REWIRE expression vectors, where we first cloned human APOBEC3A gene into pCI-Neo and then inserted different versions of PUF domains. NLS and MTS peptide sequences were synthesized (GENEWIZ) and inserted into the CU-REWIRE 1.0 vector *via* Gibson Assembly (New England Biolabs).

For the upgraded REWIREs, we assembled ten synthesized PUF repeats using PCR to generate PUF-10R (35), and replaced the PUF-8R in the original REWIRE expression vectors, producing the AI-REWIRE3.0, AI-REWIRE3.1 and CU-REWIRE2.0. For AI-REWIRE4.0, AI-REWIRE4.1 and CU-REWIRE3.0, we further modified the PUF-10R by inserting a short peptide loop (MNDGPHS) between the ninth and the tenth repeat, forming PUF-10R*.

For the wide type and mutant EGFP reporters, the coding sequences were amplified from pEGFP-C1 (Clontech) with PCR and cloned into the pCDH-CMV-MCS-EF1-Puro vector (Promega). For the reporters containing disease-relevant mutations, the coding sequences of target genes bearing pathogenic G>A or T>C mutations (as defined in ClinVar Database) were synthesized and cloned into the pCDH-CMV-MCS-EF1-Puro vector.

To clone various PUF domains that targeting different mRNAs into the REWIRE vectors, we used MultiF Seamless Assembly Mix kit (ABclonal). DNA fragments for assembly were amplified with Q5 Hot Start High-Fidelity DNA Polymerase (NEB). All vector sequences were confirmed by Sanger sequencing.

All targeted sites and primers were listed in the Table S1, and all sequences of REWIRE proteins were listed in the Supplementary Note 1.

### Mammalian cell culture and transfection

HEK 293T (ATCC CRL-3216) and SH-SY5Y (ATCC CRL-2266) cells were maintained in Dulbecco’s Modified Eagle’s Medium with high glucose (HyClone) supplemented with 10% fetal bovine serum (FBS) at 37 °C with 5% CO_2_. HCT 116 (ATCC CCL-247) cells were maintained in McCoy’s 5a Medium (Gibco) supplemented with 10% FBS at 37 °C with 5% CO_2_.

To assess the editing efficiency on the exogenous reporters, the cells were plated into a 12-well plate and transfected 12 hours later (approximate 70% confluency) with different ratios of REWIRE expression vectors and reporter plasmids at a total DNA amount of 1μg per well. The resulting cells were harvested 48 hours after transfection for further analyses. To edit the endogenous mRNAs, HEK 293T cells were transfected with 1 μg REWIRE expression plasmids and harvested 48 hours post-transfection for subsequent analyses. All transfections were performed with Lipofectamine 3000 (Thermo Fisher Scientific) in 12-well plates (or 6-well plates with adjusted plasmid amount). To control for nonspecific transfection artifacts, we also treated the cells with only transfection reagent as a mock group.

### Flp-In cell line construction

DNA fragment encoding dead EGFP (K163X) was amplified from pCDH-EGFP(K163X), and then inserted into modified pcDNA5/FRT/TO (without CMV enhancer) by Gibson Assembly. The resulting plasmid, pcDNA5/FRT/TO-EGFP(K163X), were co-transfected with pOG44 into Flp-In T-Rex 293 cells at a 1:9 ratio. 48 hours after the transfection, the cells were washed and split into fresh medium (<25% confluency), further cultured at 37°C overnight. Subsequently the cells were feed with fresh medium containing 100µg/ml Hygromycin every 3 days until the resistant colonies were visible. 3-4 individual colonies were picked for further use. The inserted sequences were also confirmed by Sanger sequencing. The expression of the target gene could be induced by adding doxycycline to a final concentration of 1 µg/ml. After 24h induction, the cells could be used to perform transfection and then be collected 2 days later for measurement of RNA editing.

### Vector construction and production of AAV

To construct the AAV vectors of REWIREs, the backbone plasmid pAV-FH vector (Vigenebio, AV88001) was linearized with Kpn1 and EcoRV, and the PCR-amplified AI-REWIRE 4.1 and CU-REWIRE 3.0 were then cloned into linearized vector by Gibson Assembly. The resulting plasmid was named as pAV-AI-REWIRE 4.1 or pAV-CU-REWIRE 3.0. The AAV9 particles were produced by Vigenebio using HEK 293T cells using the triple-transfection protocols and purified with iodixanol gradient with standard protocols.

### Flow cytometry

To assess the correction of K163X mutation in EGFP, the transfected cells were analyzed using flow cytometry at indicated time point with CytoFLEX S (Beckman Coulter). The percent of EGFP-positive cells and fluorescence intensity were calculated to reflect editing efficiency. The data was analyzed and visualized by CytExpert software (Beckman Coulter).

### Western blot

The HEK 293T cells were lysed in 1× SDS-PAGE loading buffer (Beyotime) and heated at 95°C for 10 min. The mixtures were subsequently separated by 4–20% SDS-PAGE Gel (GenScript) and transferred to poly-vinylidene fluoride (PVDF) membrane. The primary antibodies, anti-FLAG antibody (F1084, Sigma-Aldrich) and anti-GAPDH antibody (14C10, CST), were used at a 1:2000 dilution as per manufacturer’s instructions. For the secondary antibody, the HRP-linked goat anti-mouse IgG (CST#7076) and goat anti-rabbit IgG (CST#7074) were used at a 1:5000 dilution. The HRP conjugated secondary antibodies were visualized by using an enhanced chemiluminescence detection kit and ChemiDoc Touch Imaging System (Image Lab, Bio-Rad).

### RNA extraction and reverse transcription

The RNAs were extracted using the TRIzol (Thermo Fisher) following the manufacturer’s instructions. 1 μg RNA was reverse transcribed using the PrimeScript RT reagent Kit with gDNA Eraser (Takara). The cDNA was then amplified for further analyses using deep sequencing on Illumina NextSeq platform. Extraction of genomic DNA was performed using the Mammalian Genomic DNA Extraction Kit (Cat#D0061, Beyotime).

### Animal experiments

All experiments with live animals were approved by the Institutional Animal Care and Use Committee (IACUC) of the Shanghai Institute of Nutrition and Health, Chinese Academy of Sciences. B6-EGFP mice (C57BL6-Tg (CAG-EGFP)10sb/J) were purchased from Jackson Laboratory. The male and female mice were evenly assigned to groups of predetermined sample size. AAV particles encoding AI-REWIRE 4.1 or CU-REWIRE 3.0 were injected into the B6-EGFP mice through tail intravenous injection (2 × 10^12^ vector genomes per mouse in 4-week-old B6-EGFP mice (5, 6)). Four weeks after injection, we collected muscles and heart samples from B6-EGFP mice and evaluated corresponding editing efficiency *via* RNA-Seq and Sanger-seq. No visible symptoms, such as a rough hair coat and moved slowly were observed in all mice after injection.

### Sanger sequencing and deep sequencing of RNA amplicons

The cDNA was amplified with gene-specific primers flanking the target sequence (all primers and next-generation sequencing amplicons are listed in Table S1). The A-to-I or C-to-U editing rates were evaluated from Sanger sequencing using the percentage of peak area at the target adenosine site according to the sequencing chromatograms. For deep sequencing of RNA amplicons, the cDNA was amplified with gene-specific primers with appropriate forward and reverse adaptor sequences. The PCR products were purified, and the amplicon libraries were constructed using the 300-cycle Mi-Seq Reagent Kit v2 or Micro Kit v2 according to the manufacturer’s protocols for paired-end sequencing (2×150bp) on the Illumina Hi-Seq X Ten machine by the Omics core facility of PICB. The RNA editing rates were evaluated by the percentage of the edited reads at the target sites (see below).

### RNA-seq experiments

The cells were harvested 48 hours after transfection, and total RNAs were purified with TRIzol Reagent (Thermo Fisher). For each sample, 1μg of total RNAs were used to prepared ribo-minus RNA-seq libraries using the KAPA Stranded RNA-Seq Kit with RiboErase (Roche, KK8484), and the depletion of rRNAs confirmed by fluorometric quantification using the Qubit RNA HS Assay kit (Invitrogen). The sequencing was performed on an Illumina Hi-Seq X Ten machine by the Omics core facility of PICB (2 × 150-base pair paired end; 50M reads for each sample). The summary of RNA-seq data was listed in Table S2.

### RNA editing analysis using RNA-seq data

For deep sequencing analysis, we first conducted quality control and trimming using TrimGalore (0.6.0). To analyze sequences near the targeted sites, the sequence index was generated using the targeted site sequence (upstream and downstream 200-nt) of each transcript. The RNA-seq reads were aligned to the index and quantified using BWA(0.7.17-r1188) (39). Alignment BAMs were then sorted by Samtools (1.9) (40), and the RNA editing sites were analyzed using REDItools (1.2) (41) with the following parameters: -t 20 -p -u -m20 -T6-6 -W -v 1 -n 0.0 -e -d -l. AWK scripts were used to filter out all the A-to-I or C-to-U conversions within the REWIRE targeting region, which were considered significant if the Fisher’s exact test yielded a p-value less than 0.05 after multiple hypothesis correction by Benjamini Hochberg correction and at least 2 of 3 biological replicates identified the edit site. The mutations that appeared in both control and experimental groups were considered as single nucleotide polymorphisms.

To analyze entire transcriptome, AWK scripts were used to filter out the overexpressed REWIRE gene. After trimming, the reads longer than 90-nt were mapped to the reference genome (GRCh38/hg38 for human and GRCm38/mm10 for mouse) by STAR(2.4.2) (42). We then used Samtools (1.9) to sort alignment BAMs and remove duplicated reads by using Picard MarkDuplicates.jar(2.20.1-0). RNA base-editing variants were called using REDItools(1.2) with the following parameters: -t 20 -p -P -u -U -a6-6 -A6-6 -v 1 -n 0.0 -N 0.0 -e -E -d -D -l -L. With consideration of strand information, all the A-to-I or C-to-U conversions were selected using AWK scripts. From all the called variants, downstream analyses focused solely on canonical chromosomes (1-22, X, Y and M for human, 1-19, X, Y and M for mouse).

An additional layer of filter for known variants was performed using SNPs data from dbSNP version 154 and Mouse Genome Project version 7 for human and mouse respectively. The variants shared by at least 2 out of 3 biological replicates in each sample were identified as the RNA editing sites. The RNA editing level of the mock group was treated as the background, and the global off-targets of REWIRE were calculated by subtracting the background variants. The thresholds we used to filter the SNVs are based on minimum coverage (20 reads), number of supporting reads (at least five mutated reads), allelic fraction (≥0.5%), quality of the mapped reads (>30), and base quality (>30).

For manhattan plot showing the distribution of edits across the transcriptome we set a depth cutoff at 100. The sequence motifs were generated with R package ggseqlogo(0.1) (43). All plots were generated with ggplot2 package in R.

## Statistical analysis

To assess whether REWIRE constructs perturbs natural editing homeostasis, we analyzed the global editing sites shared by the mock group and PUF-Ctrl group. The differential RNA editing rates at native A-to-I or C-to-U editing sites were assessed using Pearson’s correlation coefficient analysis. Pearson correlations of the editing rate between the mock group and PUF-Ctrl group were calculated and annotated in Figure S2C and S2D.

## RESULTS

### Engineering AI-REWIREs for specific A-to-I transformation

To engineer the artificial enzymes that specifically catalyze A-to-I editing, we fused the PUF domain with the deaminase domain of ADAR1 or ADAR2 in different orders, generating PUF-ADAR and ADAR-PUF fusion proteins (Figure S1C). We engineered the PUF domain to target EGFP (Enhanced Green Fluorescent Protein) mRNA and co-expressed them in HEK293T cells, where the A-to-I editing events were measured by sequencing the RT-PCR products. Only the PUF-ADAR configuration showed detectable base editing activity near the PUF binding site (Figure S1C), and thus this configuration was used for the future design of all the AI-REWIREs (Figure 1A).

**Figure 1.**
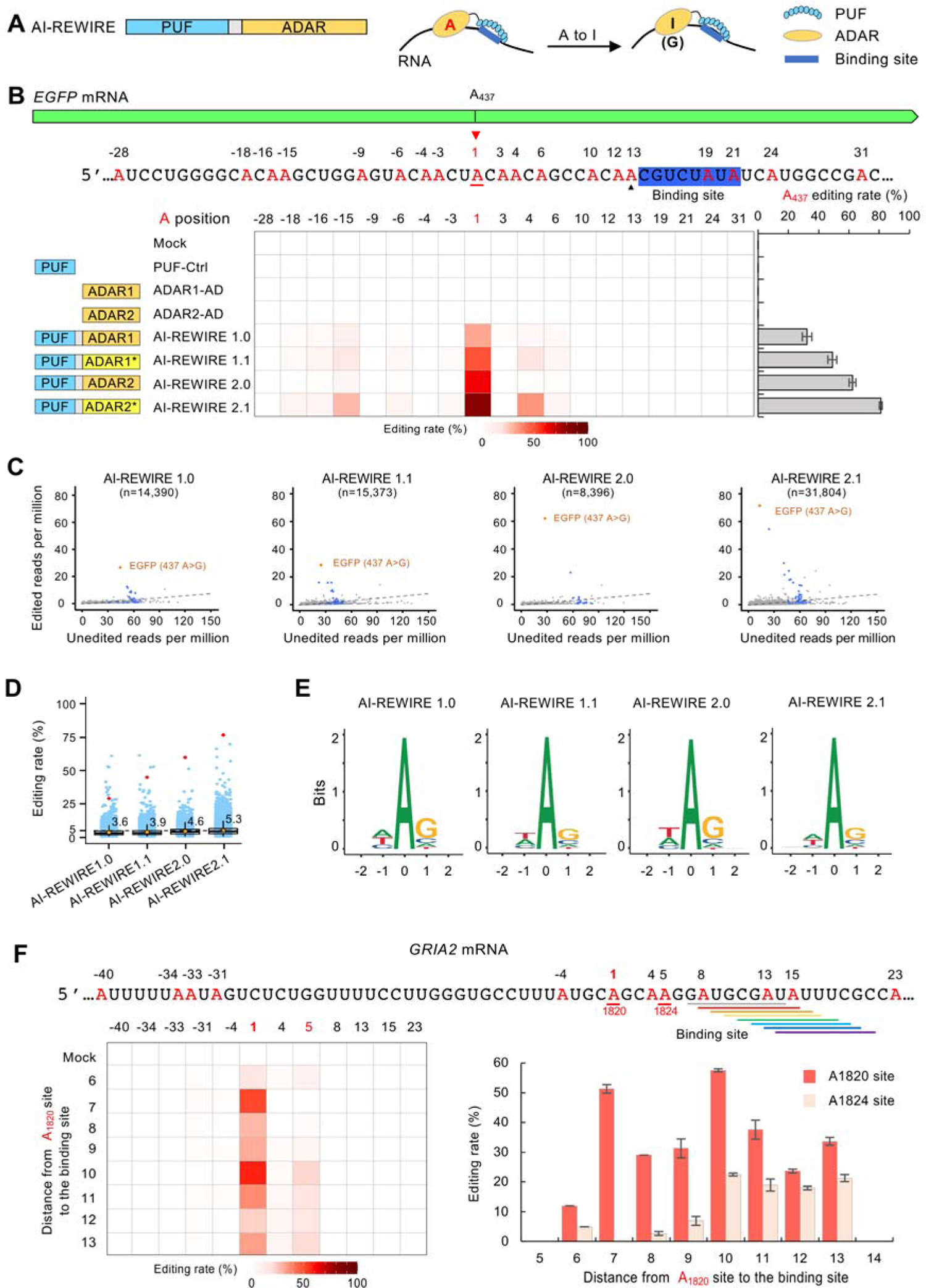
Engineering AI-REWIREs for A-to-I transformation at specific positions. (A) Engineering AI-REWIREs for specifically A-to-I editing. Domain configuration of AI-REWIRE (left); schematic of AI-REWIRE mediated specific A-to-I base editing on RNAs (right). (B) Efficient editing of EGFP mRNA by different versions of AI-REWIREs. The PUF binding site in EGFP is highlighted in blue, with the nearby adenosines marked in red (top). Heatmap showed the editing rates of all adenosines near the on-target site A437 (position 1) for each AI-REWIRE and control. The on-target editing rate of A437 was shown at right. The editing rates presented were measured by RNA-seq with triplicates (see methods). (C) Scatter plots of transcriptome-wide A-to-I RNA editing in the samples treated with AI-REWIREs. For each edited adenosine, the numbers of unedited reads (x-axis) and edited reads (y-axis) were plotted. The on-target site A437 is highlighted in orange, and the adenosines near the targeted site (in a 400-nt window) are labelled in blue. n, numbers of all edited adenosines detected. The 5% editing rate was indicated with dashed lines. Three independent experiments were performed (see methods). (D) Editing rates of transcriptome-wide A-to-I RNA editing in the samples treated with AI-REWIREs (data from Figure 1C with minimum coverage cutoff at 100). On-target sites are represented by red rhombuses; average editing rates of off-targets in each sample are indicated with orange dots and values are nearby; the 5% editing rate was indicated with a dashed line. (E) Sequence logos derived from edited adenosines. The analyses were conducted using RNA-seq data from samples treated with different AI-REWIREs (data from Figure 1C). (F) Editing of exogenously expressed GRIA2 mRNA by AI-REWIRE1.0 that binds at different distances from the intended editing site (A1820). Left, the editing rates of all adenosines near the binding site were plotted against the distance from the editing site to the 5’ end of PUF binding site. Right, the editing rates of two major sites were plotted against the binding distance (mean ± s.e.m. n=3).

We engineered various versions of AI-REWIREs using wild type ADARs or their hyperactive mutants (ADAR1 in AI-REWIRE1.0, ADAR1-E1008Q in AI-REWIRE1.1, ADAR2 in AI-REWIRE2.0, ADAR2-E488Q in AI-REWIRE2.1) (37, 38), and found that all AI-REWIREs showed efficient base editing at the on-targeted site A437 of the EGFP mRNA with highest editing rate at ∼81% (Figure 1B and Figure S1D and S1E). In addition, the editing efficiency was also affected by the expression levels of AI-REWIREs, as judged by co-expression of AI-REWIREs and the target mRNAs in different ratios (Figure S1F). As expected, the editing sites occur at the upstream of AI-REWIRE binding region, consistent with that the PUF domain binds RNA in an antiparallel fashion (25). As a control, the expression of PUF or ADAR alone did not induce detectable editing near the targeted region (Figure 1B). The estimated editing rates are highly correlated between the measurements with Sanger sequencing and RNA-seq (Figure S2A and S2B). While the Sanger sequencing provides a more accurate picture at the targeted site and the bystander editing site near the REWIRE binding window, the RNA-seq can give a global picture for the transcriptome-wide off-target effect.

Transcriptome-wide RNA-seq revealed that the major off-target A-to-I editing sites by AI-REWIREs are in the targeted EGFP mRNA rather than endogenous RNAs (Figure 1C, blue vs. grey). As expected, the AI-REWIRE1.1 and AI-REWIRE2.1 have higher editing activities at both the target site and the off-target sites compared to those with wild-type domains (version 1.0 and 2.0) (Figure 1C and 1D). In addition, we found no significant difference in global editing of endogenous sites between cells transfected with PUF domain only and mock vector (Figure S2C and S2D), suggesting PUF domain had little impact on normal editing by endogenous ADARs. Consistent with sequence preferences of ADARs (44), the AI-REWIREs were found to have a slight preference of guanine next to the edited adenosines (Figure 1E).

To determine the optimal distance between the PUF binding site and the editing site, we constructed several AI-REWIREs that specifically bind to the shifting positions at downstream of a natural edited site in GRIA2 mRNA (Figure 1F and Figure S3A). We observed an optimal editing window between the position 7 to 9 from the PUF binding site to the on-target editing site and with a low bystander editing (Figure 1F). When the binding sites of AI-REWIREs were shifted toward 3’ end, editing of a downstream adenosine (A1824) was increased (Figure 1F and Figure S3A). Our subsequent results suggest that AI-REWIREs had a fairly narrow window for optimal editing (∼3 nt) (Figure S3B and S3C).

### Application of AI-REWIREs in base correction

We next applied this system to repair a nonsense mutation (K163X) in EGFP gene. Initially the dead EGFP was co-expressed with two versions of AI-REWIREs designed to recognize a downstream sequence CUUCAAGA, and we found that the adenosine was efficiently edited to restore the green fluorescence (Figure S4A), enabling a functional rescue of EGFP in 40-60% transfected cells as judged by flow cytometry (Figure S4B). Furthermore, we generated a stable cell line containing a single copy of dead EGFP stably integrated into the genome by Flp-In system (Figure 2A). The dead EGFP was expressed from a moderate CMV promoter without the proximal enhancer to mimic the expression level of a typical endogenous gene. We expressed the two versions of AI-REWIREs in different amounts (Figure 2B), and found a dose-dependent base editing of K163X site, reaching 20% editing rate with the AI-REWIRE2.1 (Figure 2C-2E). We also monitored the phenotypic change for several days after AI-REWIRE transfection and found that the EGFP signals appeared at day one after transfection, reached the peak at day 2, and faded out after day 4 (Figure 2F).

**Figure 2.**
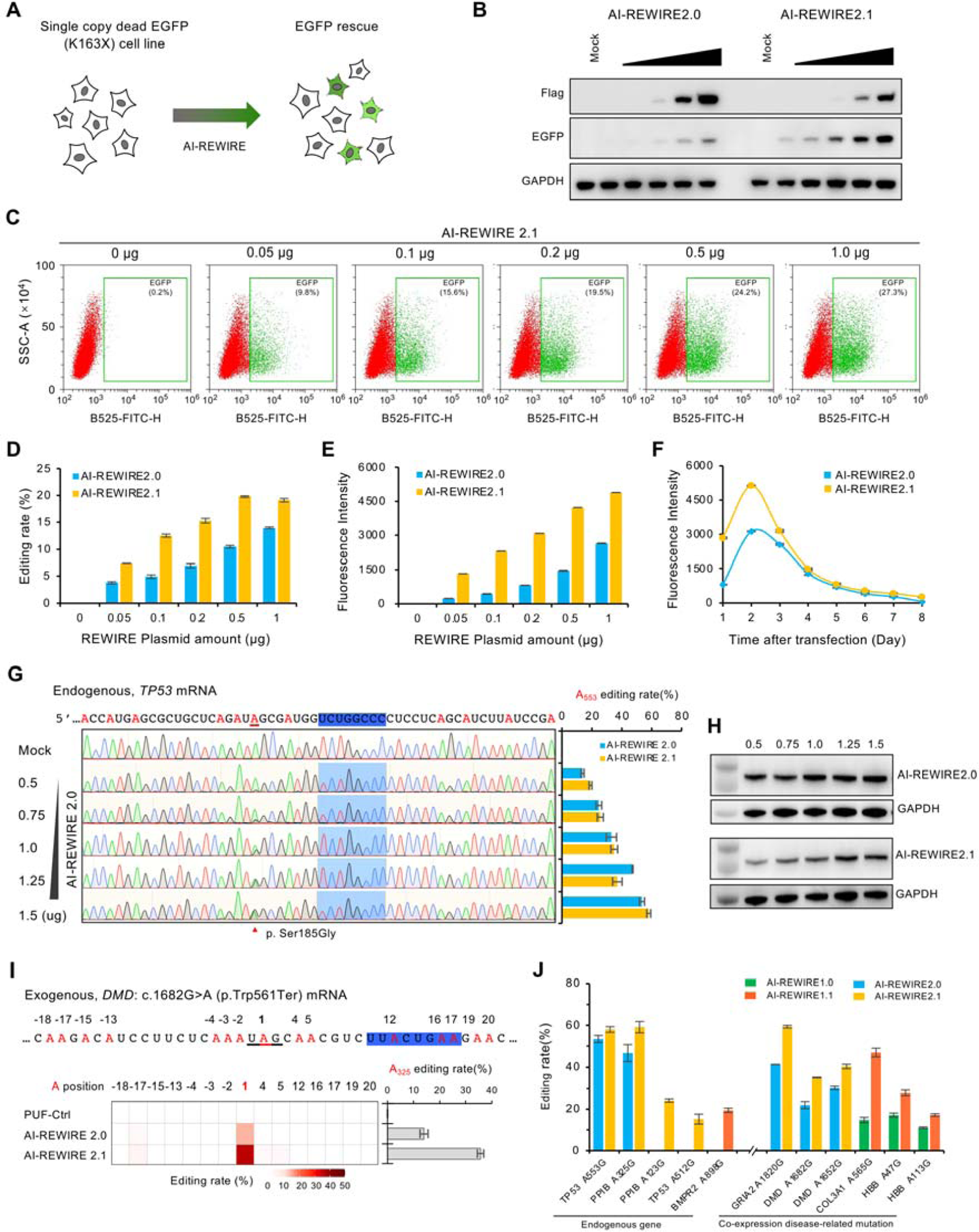
Application of AI-REWIREs. (A) Using AI-REWIREs to rescue dead EGFP. The K163X mutation was introduced to create a premature stop codon in the EGFP gene that was stably integrated into 293 cells with Flp-In system. The A-to-I editing on this site can restore the translation of full-length EGFP. (B) Western blot of expression level of AI-REWIREs and the rescued EGFP, with detection of GAPDH as loading controls. (C) Flow cytometry of dead EGFP cell lines transfected with different amounts of AI-REWIRE2.1 using a 12-well plate. The cells were analyzed 48h post transfection. (D) Editing rates of the K163X (measured by Sanger sequencing) in dead EGFP cells. The experiment conditions were same as panel C. (E) Fluorescence intensity of the rescued dead-EGFP cells. The experiment conditions were same as panel C. (F) Duration of RNA editing by the AI-REWIRE2.0/2.1, judged by the fluorescence intensity at each day after a single transfection of 1 μg plasmid. (G) AI-REWIREs edited the TP53 A553 site in a dose-dependent manner (see methods). The editing rates of all adenosines nearby were measured by Sanger sequencing (Left), with the editing rate of the on-target A553 site shown in right. Values represent mean ± s.e.m. n=3. (H) Western blot of expression level of AI-REWIREs in different co-expression experiments (samples are consistent with Figure 2G). The AI-REWIERs were Flag-tagged and the GAPDH were used as a loading control. The western blot was done in technical duplicate. (I) A-to-I editing of DMD 1682G>A mutation that is associated with Duchenne muscular dystrophy disease, the Deep-seq data showing the 1682G>A in DMD is corrected to varying levels with different AI-REWIREs. (J) Editing of exogenous disease-related G>A mutations and endogenous sites using AI-REWIREs specifically designed to recognize the target mRNAs. The mutations were selected from ClinVar. (mean ± s.e.m. n=3).

We further designed AI-REWIREs to edited several endogenous genes, including the cancer-associated TP53 gene that was previously edited by other RNA editing tools (4, 16). Using different amounts of plasmids in transfection, we again found a dose-dependent editing of the A553 in TP53 gene with efficiency ranging from 13% to 58% with two versions of AI-REWIREs (Figure 2G and 2H). For the five endogenous editing sites tested, we achieved 15-59% editing using different AI-REWIREs (Figure 2J and Figure S4C), suggesting that AI-REWIREs are capable of editing endogenous mRNAs although the editing efficiency remains to be improved.

We further examined whether AI-REWIREs could effectively edit disease-related mutations in human cells. Different versions of AI-REWIREs were engineered to target several mutations, including two DMD mutations that cause Duchenne muscular dystrophy. We co-expressed the mutated genes with corresponding AI-REWIREs, and found that the mutated base can be corrected efficiently, with AI-REWIREs yielding higher editing rates ranging from 20% to 60% (Figure 2I-2J and Figure S4D-4E).

### Engineering CU-REWIREs for specific C-to-U conversion

The successful engineering of AI-REWIREs prompted us to employ similar design for C-to-U editing. We combined the *PUF* domain and the cytosine deaminase *APOBEC3A* (apolipoprotein B mRNA editing enzyme catalytic subunit 3A, or A3A) with a peptide linker to generate the CU-REWIRE (Figure 3A). As a proof of concept, we tested the activity of CU-REWIREs on *EGFP* mRNA and found that the CU-REWIRE could enable precise C-to-U editing with efficiency up to ∼60% (Figure 3B and Figure S5A). Additionally, the CU-REWIRE(Mut) containing an inactive A3A mutation (E72A) did not show any detectable editing (45), suggesting that the base editing is indeed catalyzed by A3A (Figure 3B).

**Figure 3.**
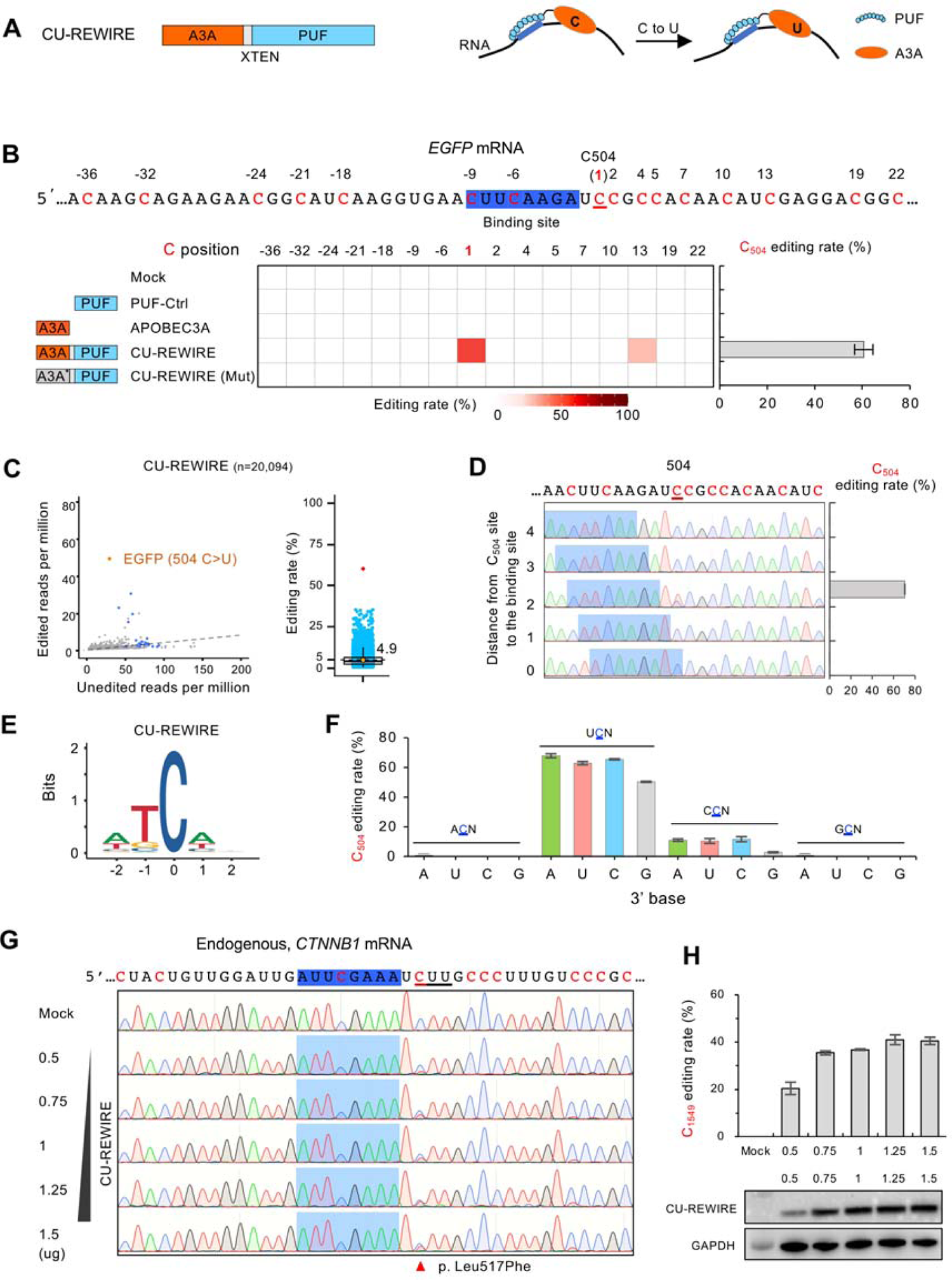
Engineering CU-REWIREs for targeted C-to-U conversion. (A) Engineering CU-REWIRE for specific C-to-U editing. Left, Domain configuration of CU-REWIRE; right, schematic of CU-REWIRE mediated specific C-to-U base editing on RNAs. (B) Efficient editing of EGFP mRNA by CU-REWIRE. The PUF binding site in EGFP and the nearby cytidines were marked (The experiments and analyses are similar to Figure 1B), with the C504 as the on-target editing site. CU-REWIRE with mutated deaminase domain was included as negative control. (C) Transcriptome-wide C-to-U RNA editing in the sample treated with CU-REWIRE (labels and cutoff are similar with Figure 1C and 1D). Left: the on-target site C504 is highlighted in orange, and the cytidines near the targeted site are labelled in blue; right: the on-target site C504 is highlighted in red; n, numbers of all edited cytidines detected. The 5% editing rate was indicated with dashed lines. (D) On-target editing rates of CU-REWIRE with different editing distances from binding sites, which represent the number of bases between the editing site and 3’ end of the PUF binding site. (E) Sequence logos derived from edited cytidines. The analyses were conducted using RNA-seq data from samples treated with different CU-REWIRE (data from Figure 3C). (F) Editing activity of CU-REWIRE on all combinations of 3-nt flanking bases around the candidate cytidine on the EGFP transcript. (G) Editing of CTNNB1 mRNA with different doses of CU-REWIREs expression vectors. The editing rates of all cytidines nearby the PUF binding site were measured by Sanger sequencing. Values represent mean ± s.e.m. n=3. (H) Editing rates of on-target C1549 at CTNNB1 with different amounts of CU-REWIREs. The rates were measured by Sanger sequencing (top). Values represent mean ± s.e.m. n=3. The protein expression level of CU-REWIREs in different co-expression experiments with GAPDH as a loading control (bottom). The western blot was done in technical duplicate.

We further used RNA-seq to examine the off-target C-to-U editing induced by CU-REWIRE, and found some off-target editing sites in both the targeted EGFP mRNA and other endogenous transcripts (Figure 3C). Interestingly, the endogenous off-target sites with high editing rates are mostly expressed in a very low level, which might be contributed by an “editing noise” (see discussion for more details). Compared to AI-REWIREs, the CU-REWIRE appeared to have a tighter editing window and performed C-to-U editing at position 2 after the PUF binding site (Figure 3D). Moreover, the edited cytidines were predominantly found within the short consensus motif UC (Figure 3E). Such preference was experimentally validated using all base combinations around the targeted cytidine, where the UCN sequence was strongly edited by the CU-REWIRE and the CCN was also tolerated to a less extend (Figure 3F). This finding is consistent with the target preference of APOBEC3A (45). Interestingly, we found that the guanosine at immediate downstream of the cytidine can reduce editing efficiency, reflecting an unreported feature for APOBEC3A target preference (Figure 3F).

To test if CU-REWIREs are capable of editing endogenous mRNAs, we designed a CU-REWIRE to target an endogenous gene CTNNB1, and achieved 20-45% editing efficiency at the targeted C1549 site in a dose-dependent fashion (Figure 3G and 3H). To further test the general applicability in RNA base correction, we designed CU-REWIREs targeting two disease-related T>C mutations in human genes (*EZH2* in Weaver syndrome, *SCN1A* in Dravet syndrome). We found that the deleterious mutations were indeed corrected *via* cytidine editing with efficiency ranging from 21% to 29% (Figure S5B and S5C). Furthermore, we also targeted the C246 in *SARS-Cov-2* membrane protein, and achieved high editing efficiency (∼62%) (Figure S5D).

### Improvement of the REWIREs

To optimize the REWIREs, we redesigned a PUF domain with 10 tandem repeats (PUF_10R) that can specifically recognize RNAs by a 10-nt binding site rather than an 8-nt site of wild type PUF_8R, thus should significantly increase its specificity and reduce off-targets (27, 35). We applied this design strategy to generate AI-REWIRE3.0 and AI-REWIRE4.0 using two versions of PUF_10R (Figure 4A, see methods), and further engineered AI-REWIRE3.1 and AI-REWIRE4.1 by upgrading deaminase domain with hyperactive ADAR2. As expected, the AI-REWIREs with PUF_10R substantially reduced both the editing rates and the numbers of off-target sites without sacrificing the on-target editing efficiency (Figure 4B and 4C). Compared to the most active AI-REWIRE2.1 (on-target editing rate=81%, off-target counts=31804), the updated AI-REWIRE3.1 had much less off-target counts (1908) and a decent editing rate (∼62%) while the AI-REWIRE4.1 showed similar tendency (on-target editing rate=72%, off-target counts=6664) (Figure 4C and 4G). Interestingly, such improvement was achieved with a lower expression level than the early versions (Figure S6A and S6B), which is a desirable feature for *in vivo* application where the high expression of engineered editase may be difficult to achieve and/or have adverse effects. We further used the upgraded AI-REWIRE4.0 to edit an endogenous gene (PPIB), and found that it achieved a comparable editing rate to the AI-REWIRE2.0 targeting the same site (Figure S6C), but with much less off-targeting effect (Figure S6D), which is consistent with its longer recognition sequence (10nt *vs.* 8nt). Since the PUF domain recognizes a short RNA sequence, this specific AI-REWIREs has other recognition sites in the transcriptome, some of which also being edited (brown dots in Figure S6D). This is an inherent limitation of all engineered factors using PUF domain, which could be managed through target design and additional controls in protein expression (see discussion). As expected, both AI-REWIREs function in a dose-dependent manner in editing endogenous targets (Figure S6E).

**Figure 4.**
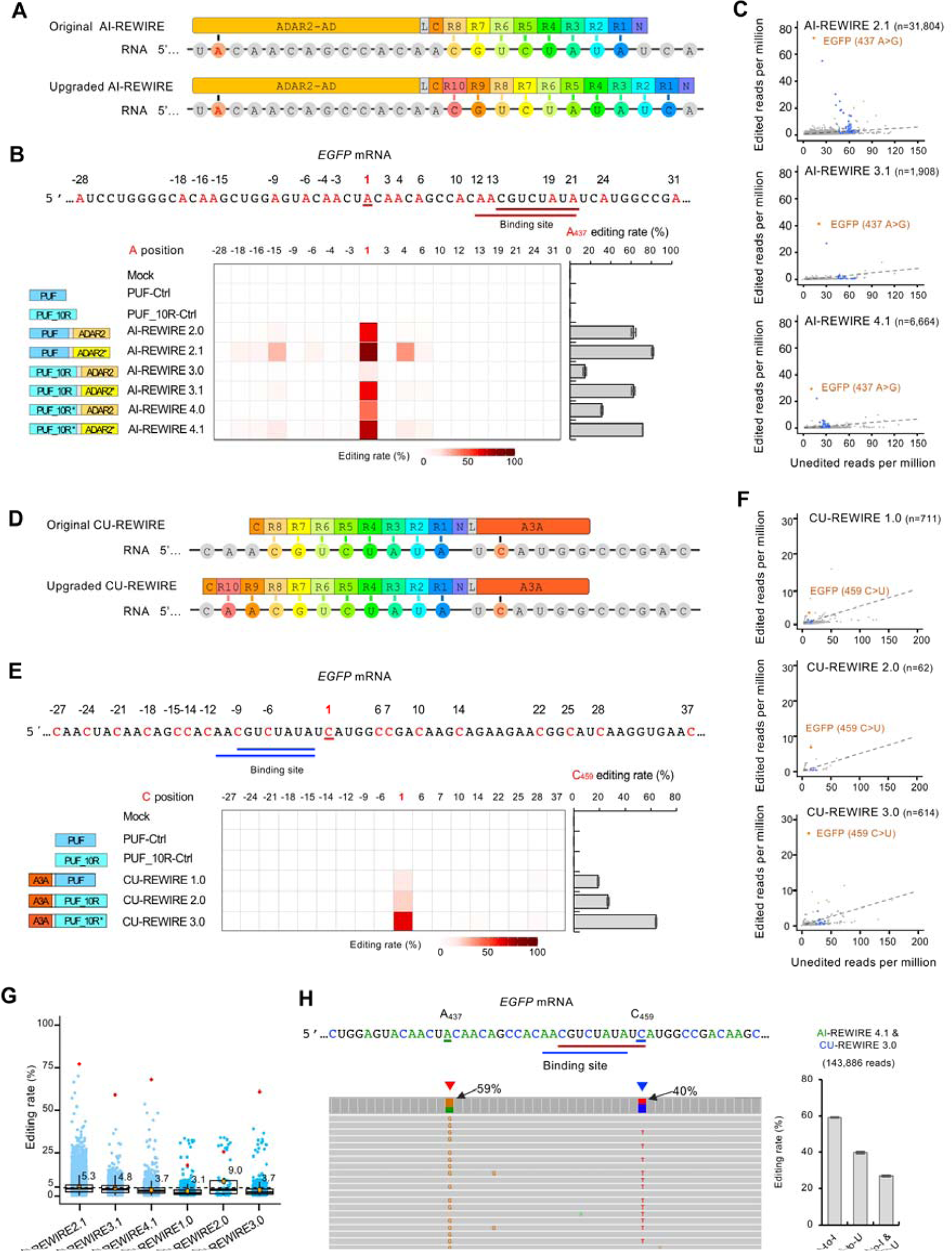
Optimization of the REWIREs. (A) Schematic of AI-REWIREs with upgraded PUF variants. Top, the original AI-REWIRE with the PUF-8R that recognize targets by 8-nt sequence. Bottom, upgraded AI-REWIRE with the PUF-10R that recognize targets by 10-nt sequence. (B) Efficient editing of EGFP mRNA by upgraded versions of AI-REWIRE. The PUF binding site in EGFP is underlined in red. The experiments and analyses were similar to Figure 1B. The AI-REWIRE 3.0, 3.1, 4.0 and 4.1 all used PUF_10R scaffold as target recognition domain, with 4.0 and 4.1 containing a short loop between PUF repeat 9 and 10 (see method). (C) Scatter plots of transcriptome-wide A-to-I RNA editing with edited/unedited reads (labelled as Figure 1C) for the samples treated with AI-REWIREs. (D) Schematic of CU-REWIREs with upgraded PUF variants. (E) Efficient editing of EGFP mRNA by upgraded versions of CU-REWIRE. The PUF binding site in EGFP is underlined in blue, with the nearby cytosines marked in red (on-target site, C459). The experiments and analyses were similar to Figure 3B. The CU-REWIRE 2.0 and 3.0 all use PUF_10R as target recognition domain (see methods). (F) Scatter plots of transcriptome-wide C-to-U RNA editing in the samples treated with CU-REWIREs (labeled as Figure 3C left panel). (G) Editing rates of transcriptome-wide RNA editing in samples treated with different REWIREs (data from Figure 4C and 4F, labels and cutoff are same as Figure 1D). (H) Successive editing of adenosine and cytidine with the AI-REWIRE 4.1 and CU-REWIRE 3.0 that bind to the same site on EGFP (the binding site is underlined in red and blue respectively). The experimental condition is similar with Figure 4B and 4E except that half amounts of each vector were transfected. The experiments and sequencing were carried out in duplicates. Left, representative Deep-seq reads surrounding the target site; right, the editing rates calculated using the number of reads supporting single and concurrent editing divided by the total reads.

With the same strategy, we upgraded the original CU-REWIRE (CU-REWIRE1.0) into CU-REWIRE2.0 and CU-REWIRE3.0 using different versions of PUF_10R (Figure 4D and 4E), and measured their editing efficiency on *EGFP* mRNA (Figure 4E and Figure S6B). While all the CU-REWIREs showed precise editing at the targeted cytidine (C459), the CU-REWIRE3.0 achieved the highest editing rate (up to 65%) and relatively low off-target effects across the entire transcriptome (Figure 4F and 4G), indicating a comprehensive improvement. We designed new CU-REWIRE3.0s targeting two disease-related T>C mutations in *EZH2* and *SCN1A* genes, and found that these mutations were indeed corrected *via* cytidine editing with modestly increased efficiency while the target specificity was improved (Figure S6F and S6G).

To take advantage of REWIREs that can introduce multiple variants in RNAs, we applied AI-REWIRE and CU-REWIRE together for concurrent editing on a single transcript, producing efficient A-to-I and C-to-U editing simultaneously (Figure 4H and Figure S7). We also noticed that using two REWIREs in combination showed comparable editing rates to the AI-REWIRE-4.1 or CU-REWIRE-3.0 alone at the single base (compare Figure 4H to Figure 4B and 4E), suggesting they may function independently. Importantly, the overlap of the binding sites for two editases did not interfere with the concurrent editing (Figure 4H), suggesting they undergo dynamic turn over on the target RNA like what would be expected for *bona fide* enzymes. Within each targeted mRNA molecule, both AI-REWIRE 4.1 and CU-REWIRE3.0 preferably edited on the single intended site as judged by reanalyzing the RNA-seq data (Figure S8). In addition, we also tested the REWIREs in additional two human cell lines (SH-SY5Y and HCT116), and found that different versions of REWIREs achieved comparable editing efficiency on the targeted sites (60%-86% for A-to-I editing, and 25–80% for C-to-U, Figure S9).

### RNA editing in mouse model with REWIREs

We further tested the *in vivo* editing efficacy of the REWIRE system using a transgenic mouse model (B6-EGFP) that stably expresses EGFP. The EGFP-targeting AI-REWIRE4.1 or CU-REWIRE3.0 were packed in AAV9 vectors, which were systematically delivered to 4-week-old mice using intravenous injections at the tail vein (Figure 5A and 5C). Because the AAV9 vectors has a high tropism for muscle and heart (46, 47), we collected the muscle and heart samples at 4 weeks post-injection, and evaluated the editing efficiency *via* Sanger sequencing and RNA-seq. At the tibialis anterior muscle of the three mice, the AI-REWIRE 4.1 achieved 27%-34% editing efficiency at the targeted A437 site of *EGFP* (Figure 5B), whereas the CU-REWIRE 3.0 targeting the C459 of the *EGFP* showed 44%-51% editing efficiency (Figure 5D). As a control, the expression of the PUF domain alone through same AAV9 vectors did not induce detectable editing (Figure 5B and 5D).

**Figure 5.**
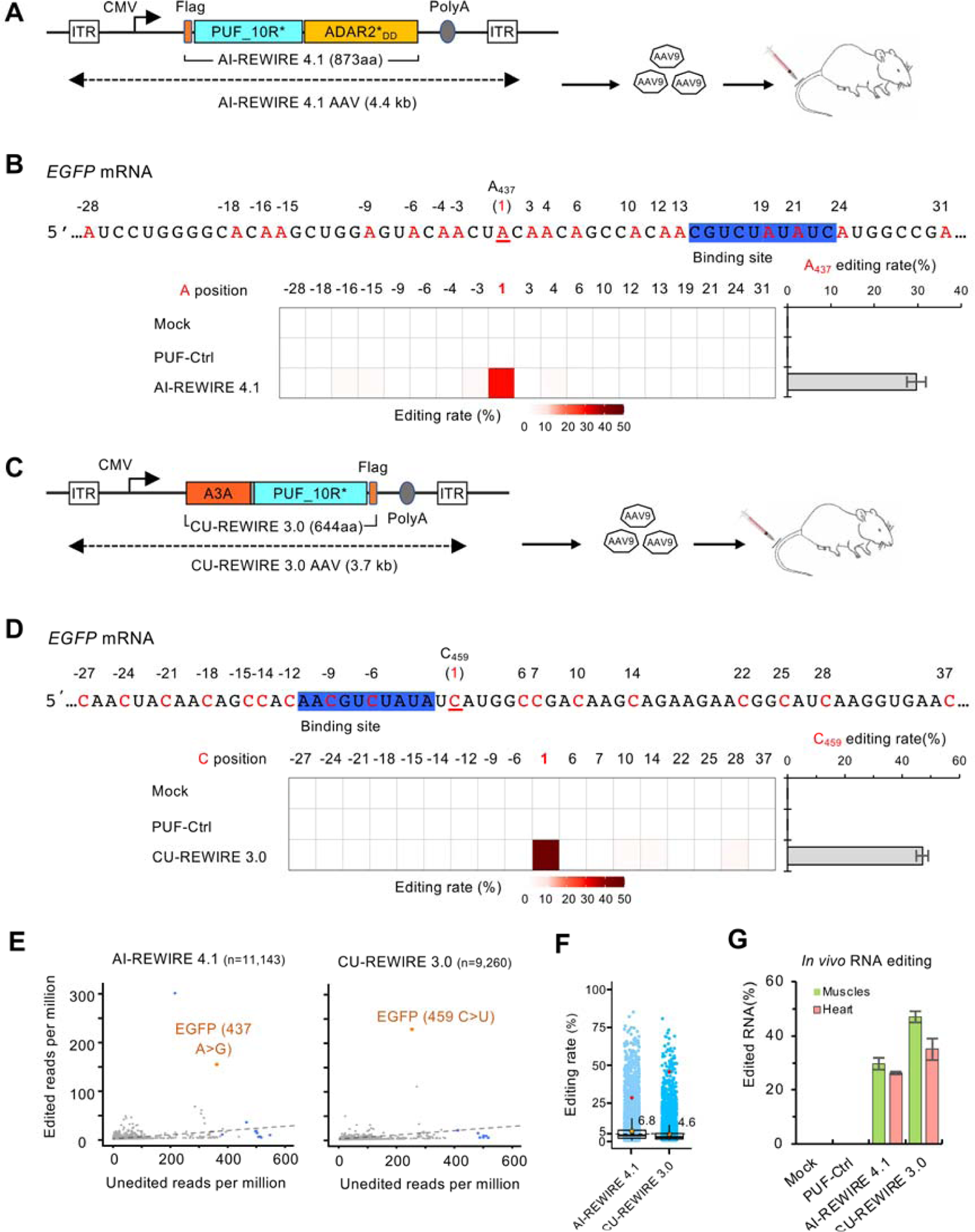
*In vivo* RNA editing in the B6-EGFP mouse model (A) Schematic for AI-REWIRE 4.1 delivered to B6-EGFP mice through tail vein IV injection of AAV9. The total insert size between the two inverted terminal repeats (ITR) was 4.4 kb. (B) Efficient Editing of EGFP mRNA by AI-REWIRE 4.1 in the tibialis anterior muscles. The PUF_10R binding site and the on-target editing site A437 (position 1) were highlighted. The editing rates were measured by RNA-seq with triplicates as described in Figure 1B. Values represent mean ± s.e.m. n = 3. (C) Schematic for CU-REWIRE 3.0 delivered to B6-EGFP mice using AAV9. The total insert size between the two ITR was 3.7 kb. (D) Efficient editing of EGFP mRNA by CU-REWIRE 3.0 in the tibialis anterior muscles. The experiments were similar to Figure 5B. Values represent mean ± s.e.m. n = 3. (E) Scatter plots of transcriptome-wide A-to-I or C-to-U RNA editing in the samples treated with respective REWIREs (labels are similar with Figure 1C and Figure 3C left panel). (F) Editing rates of transcriptome-wide RNA editing in samples treated with different REWIREs (data from Figure 5E, labels and cutoff are same as Figure 1D and Figure 3C right panel). (G) Efficient editing of EGFP mRNA in muscle and heart tissues. Values represent mean ± s.e.m. n = 3, respectively.

We further measured the low off-target editing on the mouse transcriptome using RNA-seq profiling, and found that most off-target sites were edited with low efficiency (Figure 5E and 5F). In addition, we observed effective RNA editing of the intended sites in both heart and muscles, which is consistent with the tissue preference of AAV9 (Figure 5G). These results suggested that REWIREs can effectively edit RNAs in animal, however the editing efficiency remains to be optimized.

## DISCUSSION

Compared to DNA, targeted RNA editing is reversible and more flexible, presenting a more tractable route for gene manipulation. Here we engineered a series of programmable RNA editases for specific base editing. With different functional domains, the REWIREs can effectively achieve both A-to-I and C-to-U editing, and can be used in correcting pathogenic mutations of endogenous genes. The protein sequences of this system are entirely originated from human proteome, which should avoid the innate immunity from CRISPR-based system using bacterial proteins. Importantly, the REWIRE system achieved *in vivo* base editing in mice through systematic delivery with AAV (Figure 5), suggesting its potential in personalized gene therapy.

The REWIREs consist of a single protein that recognizes target RNAs without the assembly of gRNA/protein complex, thus harboring high efficiency while avoiding assembly intermediates or misassembled by-products. The RNA-binding affinity is similar for all PUF_8Rs with similar GC content in the targets (*K_d_* in the order of nM) (25, 29), and PUF_10R can increase the binding affinity by ∼10 fold (35). A single component editase can also be transported to different subcellular compartments (e.g., mitochondria or chloroplast), which may be used in specific editing of extranuclear genes. As an RNA editing tool, the optimized REWIRE has several desirable features. First, the REWIREs with PUF_10R have a small size (873 aa and 644aa for AI- and CU-REWIREs) suitable for AAV delivery in clinical applications. Second, the REWIRE sequences are entirely originated from human genes, making it less likely to induce immune response for *in vivo* applications. Third, the REWIRE directly converts the RNA base without using endogenous repair pathways, thus may be able to edit RNA in post-mitotic cells including neurons. Finally, different REWIREs can be independently applied to achieve simultaneous A-to-I and C-to-U editing in the same transcript, allowing concurrent editing of multiple mutations that synergistically cause human diseases (Figure S8).

As a new technology, the REWIRE system also has limitations that should be improved with future optimization. First, PUF domains target a relatively short sequence (8 to 10-nt in this study), which is shorter than the CRISPR-based system but is higher than the 7-nt seed match of siRNAs. This may contribute to some unintended targets with similar recognition sites in the transcriptome. This problem could be managed by optimizing target designs. For example, the specificity could be improved by using tandem PUFs or a PUF with even more repeats (a PUF with 16 repeats has been proved feasible (35)); and computational analyses could be applied during target selection to minimize unwanted targets in transcriptome. Second, REWIREs still have bystander editing sites within the editing window, although this problem is less severe than the CRISPR-based editing tools (4, 17). It is possible to optimize the deaminase domain and the linker region to increase editing accuracy. Theoretically, we should be able to introduce mutations in the deaminase domain to increase the selectivity of editing sites, and/or modify the linker region to refine the editing window and reduce the bystander editing.

The off-target effect has been a major concern for gene editing, especially for the genomic DNA because the deleterious changes will be permanent. However, the off-target editing on RNAs may be less detrimental since the cellular RNAs constantly undergo synthesis and degradation cycles. Both the edited and unedited versions of transcripts are co-expressed in the same cells, probably damping the side effect of the off-targeting editing. The off-target effects are mainly depended on the choice of functional domains. For example, the APOBEC1 were shown to induce more off-target editing sites in both DNA and RNA than the APOBEC3A, which can be reduced by more specific deaminase (7-10,48). We found that REWIREs with ADAR2 showed less off-target effects than ADAR1, and the hyperactive mutants increased off-target effects while improving editing rates (Figure 1C and 1D). Further optimization on the PUF domain also remarkably reduced the off-target effects while maintaining the high editing efficiency (Figure 4).

In addition, the common procedures to measure editing sites using RNA-seq tend to over-estimate off-target sites, probably due to the sequencing errors associated with NGS technologies and underlying genomic variants that may confound the analyses (49). It has been reported that the accuracy for quantifying the RNA editing events can be generally improved by a higher sequencing depth and more uniform read coverage (50). Therefore, for a more reliable estimation, we sequenced the total RNAs (rather than mRNAs) with a deep coverage (∼50 million 150 bp paired-end reads for each sample) and conducted technical triplicates to reduce sequencing errors. We found that many “editing” sites covered only by a small number of reads have surprisingly high editing rates. However, such estimation may not be reliable, as increasing the cut-off for the read numbers will eliminate these “editing” sites whereas the true positive editing events remain unchanged (Figure S10). This “editing noise” is similar to a previous report that virtually all adenosines within human Alu regions could be edited as judged by an ultra-deep sequencing of selected Alu elements (>5000 reads/site) (51).

Compared to other gRNA-based RNA editing tools, including the new CRISPR-based REPAIRv2 (4) and REPAIRx (18) systems, the AI-REWIREs showed a higher editing efficiency at the targeted sites and a slightly higher off-target effect, the CU-REWIREs presented comparable properties on editing efficiency and off-target effect to the latest C-to-U editing tools, CURE systems in HEK 293T cells (19). This is a rough comparison because of the technical challenges in estimating off-target effect (Figure S10). Nevertheless, REWIRE proteins present a simple and human originated system with comparable performance. Multiple technologies may complement to each other in editing different targets, where the users should consider a tradeoff between editing efficiency and the off-target effects to find a sweet spot.

Although we have improved the REWIREs with different versions of PUFs, additional research is certainly needed to further optimize this system. Compared to previous tools using adenosine deaminase domain of ADARs for A-to-I editing and modified ADAR2 for C-to-U editing (4), (17), we achieved C-to-U editing with cytidine deaminase Apobec3A that is smaller than ADARs. The APOBAC family has many members with various specificities and strengths, therefore further exploration on the APOBAC family with possible modifications may expand the functional modules of REWIREs to edit additional bases. In addition, adjusting the composition and length of the internal linker in REWIRE may also improve its performance. Collectively, the REWIRE represents a simple and practical alternative to CRISPR-based approaches, expanding the current RNA editing toolbox.

## AVAILABILITY

The original RNA and DNA sequencing datasets have been deposited in the NCBI Gene Expression Omnibus (accession code GSE155734) and the National Omics Data Encyclopedia (accession code OEP001013) for public downloading. Custom computer code is available upon request.

## SUPPLEMENTARY DATA

Supplementary Data are available at NAR online.

## Supporting information

Supplementary Note 1. REWIREs plasmid protein sequences

## ACKNOWLEDGEMENT

We thank Ms. Yun Jiang for helps in preparing the paper, and members of Wang lab for discussion and comments of this manuscript. We thank Dr. Li Yang for the generous gift of human APOBEC3A.

## Supplementary Figures

**Figure S1.**
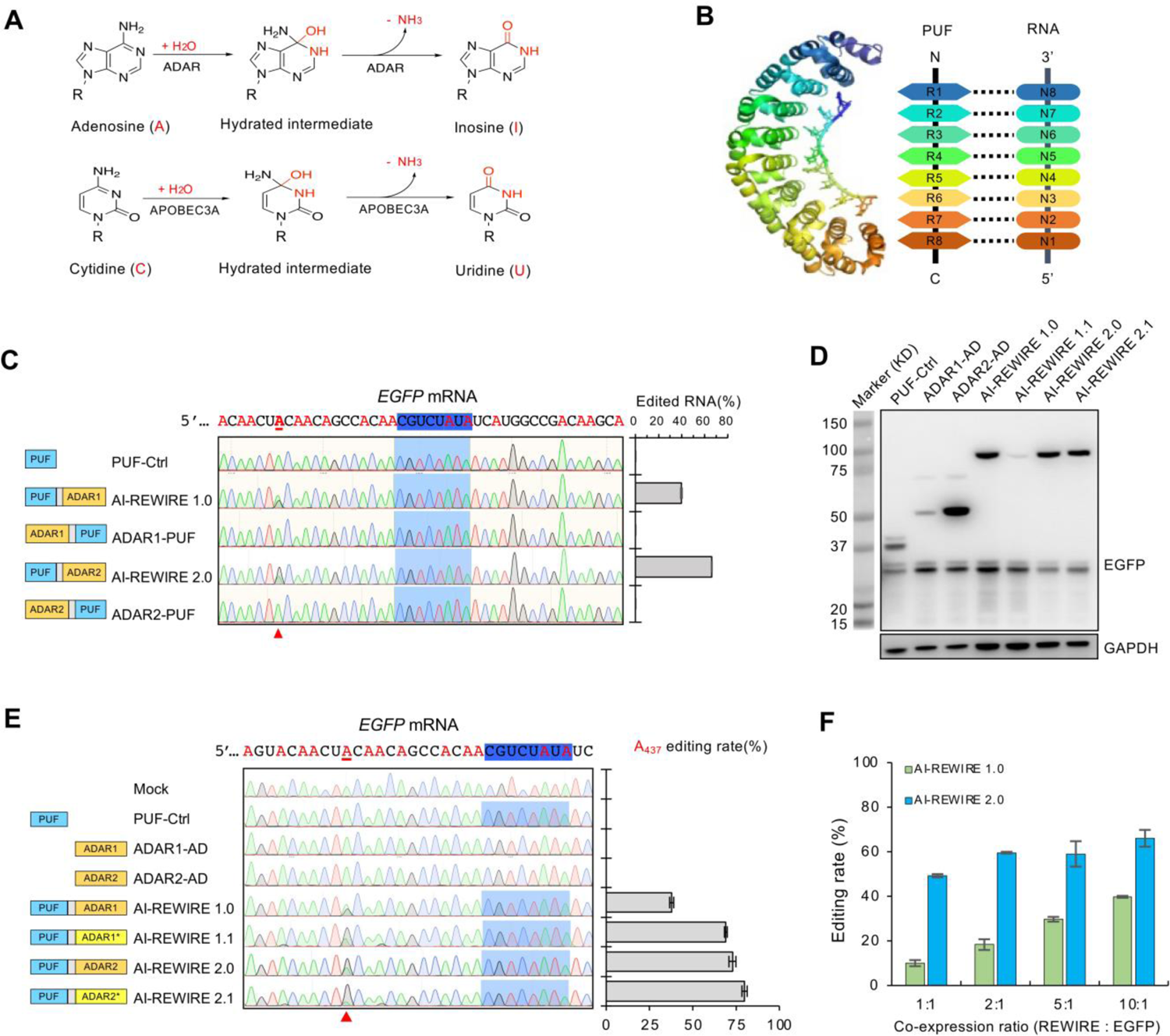
The design principle of REWIREs and assessment of specific A-to-I editing by AI-REWIREs. (A) Top, the adenosine (A) to inosine (I) editing reaction mediated by ADARs *in vivo*, where inosine will be recognized as guanosine(G) by DNA polymerases. Bottom, the cytidine (C) to uridine (U) editing reaction mediated by APOBEC3A. R, ribose in RNA. (B) Crystal structure of PUF domain (PDB: 2YJY) and the diagram of PUF-RNA interaction. The PUF domain can directly recognize its targeted RNA sequence without the co-folding and assembly steps into a gRNA-protein complex of CRISPR-Cas system. (C) Efficient editing of EGFP transcripts by various configuration combinations of PUF-ADAR and ADAR-PUF fusion proteins. Base editing rates were measured by Sanger sequencing. (D) Bar plots showing the on-target RNA base editing rates using different ratio of targets and AI-REWIREs (see methods). Base editing rates were measured by Sanger sequencing. Values represent mean ± s.e.m. (n=3). (E) Protein levels of AI-REWIREs. Representative images of western blot were from the same samples in Figure 1B, GAPDH was included as a loading control. The western blot was performed with technical duplicate. (F) Efficient editing of EGFP transcripts by various AI-REWIRE versions. For each version, the editing rate of any adenosine near the target site was measured by Sanger sequencing (left panel). The editing rate of the on-target site A437 was shown in the right panel. Values represent mean ± s.e.m. (n=3).

**Figure S2.**
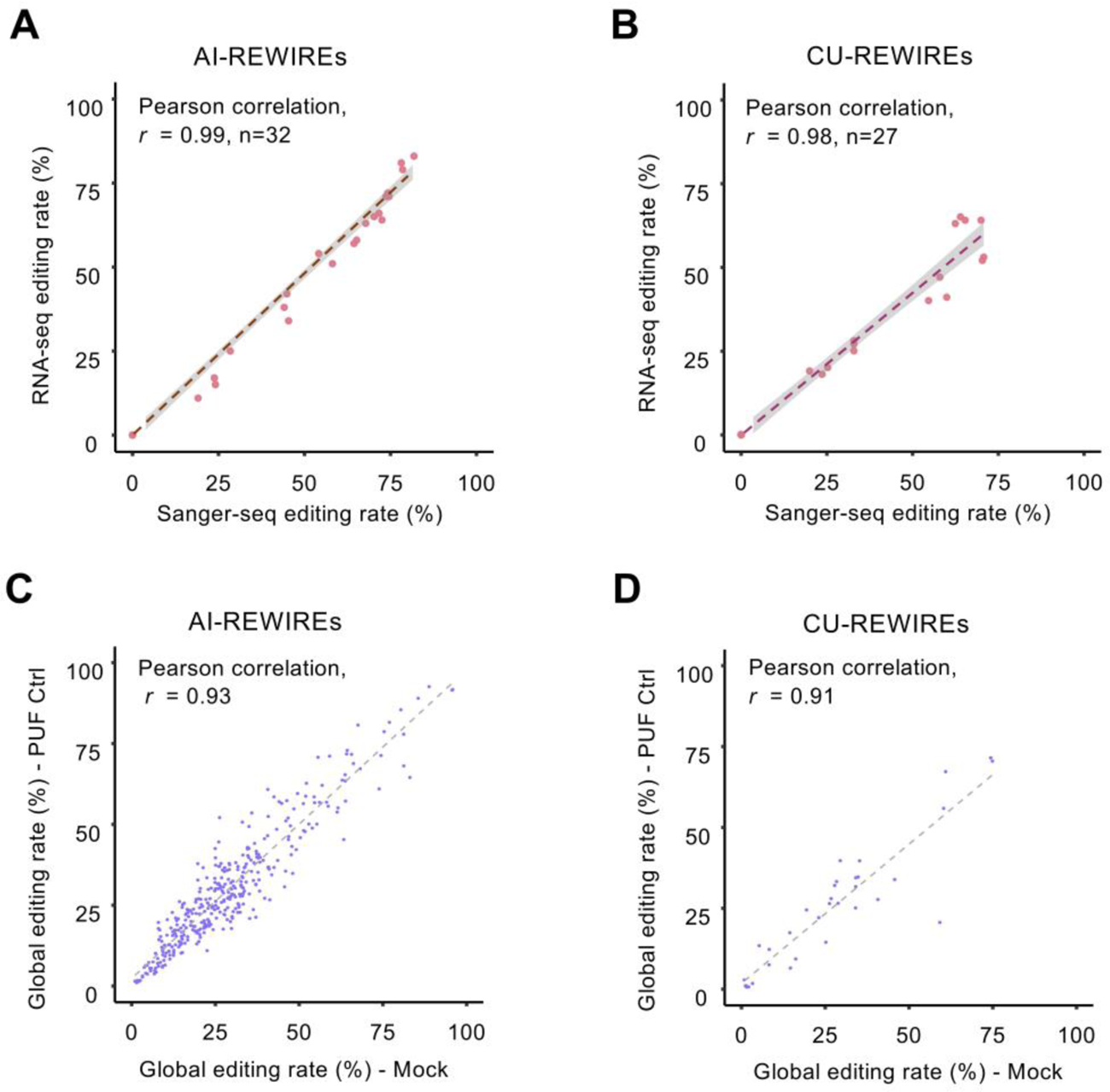
Sanger-seq and RNA-seq measurement correlation and transcriptome-wide off-target analysis of control RNA. (A) Editing rate of Sanger-seq and RNA-seq measurement correlation on the target site edited by AI-REWIREs. Each dot, an individual A-to-I editing sample targeting EGFP mRNA used in this study; x-axis, on-target editing rate measured by Sanger-seq; y-axis, on-target editing rate measured by RNA-seq. (B) Editing rate of Sanger-seq and RNA-seq measurement correlation on the target site edited by CU-REWIREs. Each dot, an individual C-to-U editing samples targeting EGFP mRNA used in this study; x-axis, on-target editing rate measured by Sanger-seq; y-axis, on-target editing rate measured by RNA-seq. (C) Transcriptome-wide off-target effects analysis of Mock control RNA and PUF-Ctrl RNA on native A-to-I editing sites. x-axis, editing rate in Mock sample; y-axis, editing rate in PUF-Ctrl sample. Three independent experiments were performed (see methods). (D) Transcriptome-wide off-target effects analysis of Mock control RNA and PUF-Ctrl RNA on native C-to-U editing sites. x-axis, editing rate in Mock sample; y-axis, editing rate in PUF-Ctrl sample. Three independent experiments were performed.

**Figure S3.**
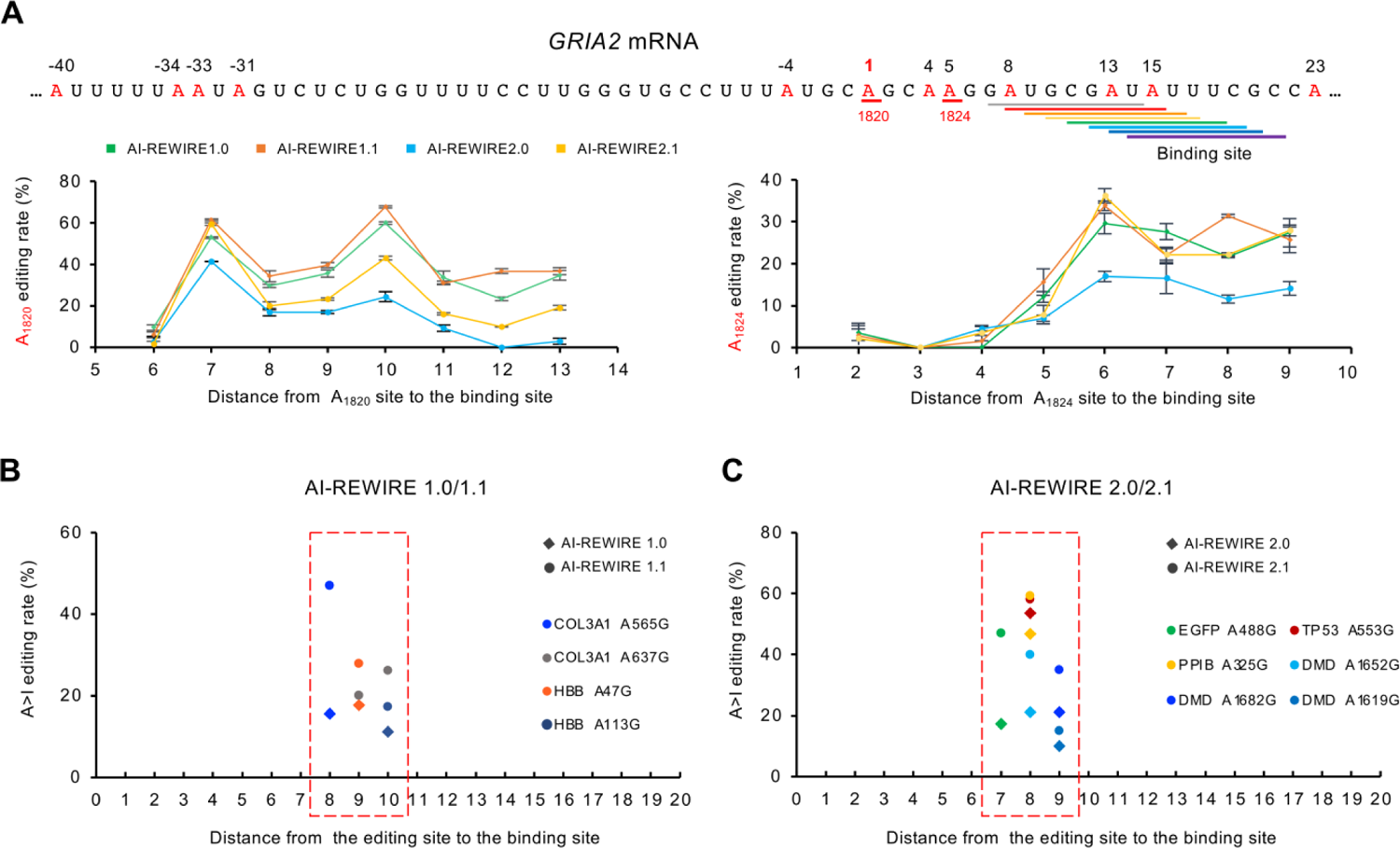
AI-REWIREs showed a narrow editing window. (A) Designed editing on GRIA2 transcripts by AI-REWIREs with step-by-step strategy to test the optimal editing window. PUF binding sites in GRIA2 mRNA were underlined in different colors, with the nearby adenosines marked in red. Left, editing rates of GRIA2 A1820 site by different AI-REWIREs that binds with shifting distances from the A1820 site. Right, editing rates of GRIA2 A1824 site by the corresponding AI-REWIREs. For each version of AI-REWIRE, the editing rate of adenosines near the targeted site were measured by Sanger sequencing. Values represent mean ± s.e.m. (n=3). (B) Summary of adenosines editing window mediated by AI-REWIRE1.0/1.1 around the target site. x-axis, distance from the editing site to the binding site, 1-based; y-axis, A-to-I editing rate. (C) Summary of adenosines editing window mediated by AI-REWIRE2.0/2.1 around the target site. x-axis, distance from the editing site to the binding site, 1-based; y-axis, A-to-I editing rate.

**Figure S4.**
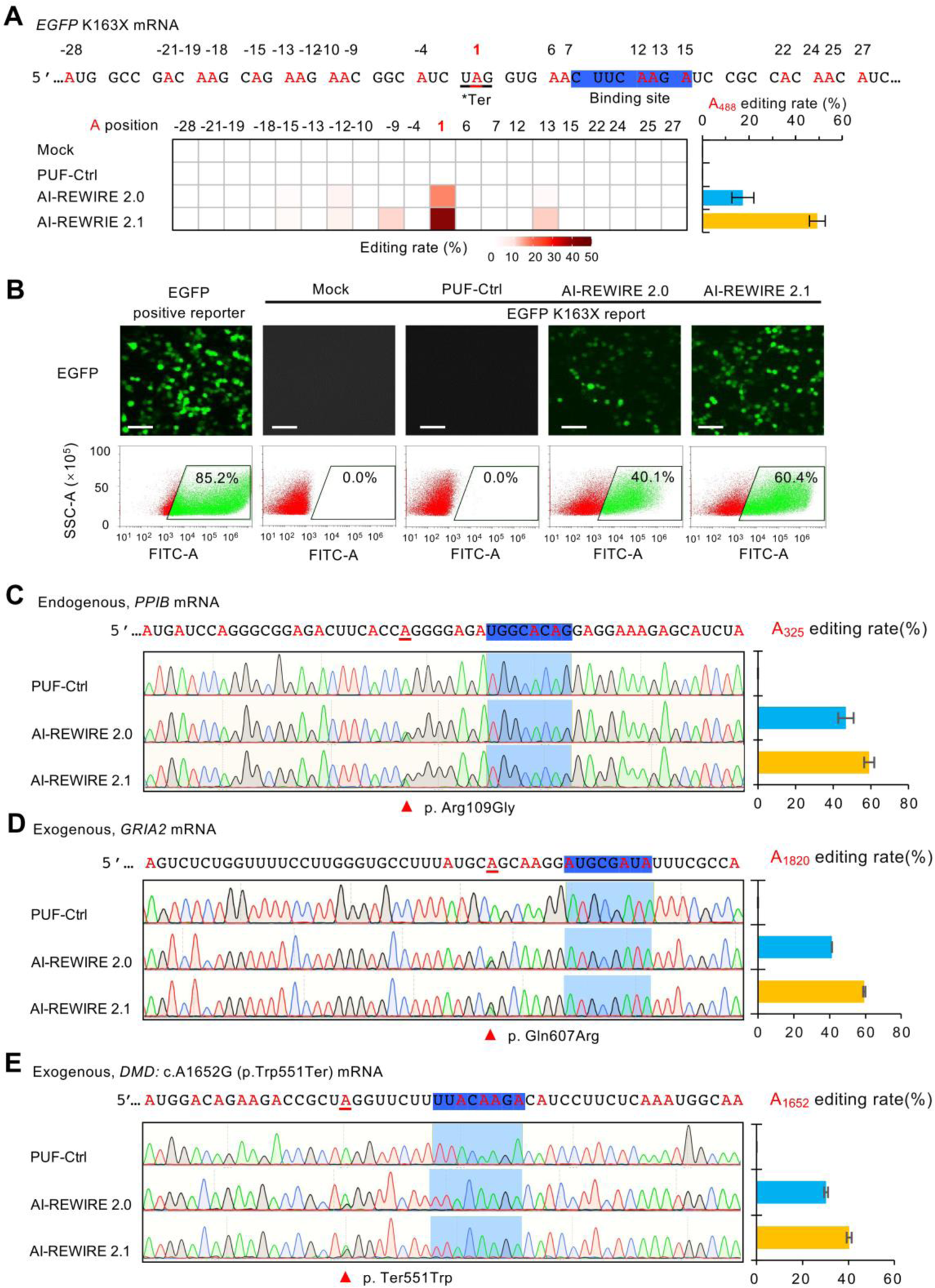
Application of AI-REWIREs on editing endogenous and exogenous pathogenetic mutant RNA. (A) Top, engineering AI-REWIREs to correct K163X mutation in EGFP with transient co-transfection. The K163X codon mutation was introduced in EGFP to cause premature termination, and the editing of this site will restore functional protein. The PUF binding site is highlighted in blue. Bottom, for each AI-REWIRE version, the editing rate of all adenosines near the target site was measured by Deep-seq (left) and the editing rate of the on-target site A437 was shown in right. Values represent mean ± s.e.m. (n=3). (B) Restoration of EGFP fluorescence by the AI-REWIREs in same samples as judged by microscopy (top) and flow cytometry (bottom). Scale bar, 50 μm. Values represent mean ± s.e.m. (n=3). (C) Editing rates for an endogenous gene, *PPIB*, with different AI-REWIREs. The editing rates of all adenosines nearby were measured by Sanger sequencing, with the editing rate of on-target A325 shown in right. Values represent mean ± s.e.m. (n=3). (D) Engineering AI-REWIREs to edit *GRIA2* A1820 site that is associated with ligand-gated ion channel in the central nervous system. The editing rates of all adenosines nearby were measured by Sanger sequencing (left), with the editing rate of on-target A1820 shown on the right. Values represent mean ± s.e.m. (n=3). (E) Engineering AI-REWIREs to correct *DMD* 1652G>A mutation that is associated with Duchenne muscular dystrophy disease. The editing rates of all adenosines nearby were measured by Sanger sequencing (left), with the editing rate of on-target A1652 shown on the right. Values represent mean ± s.e.m. (n=3).

**Figure S5.**
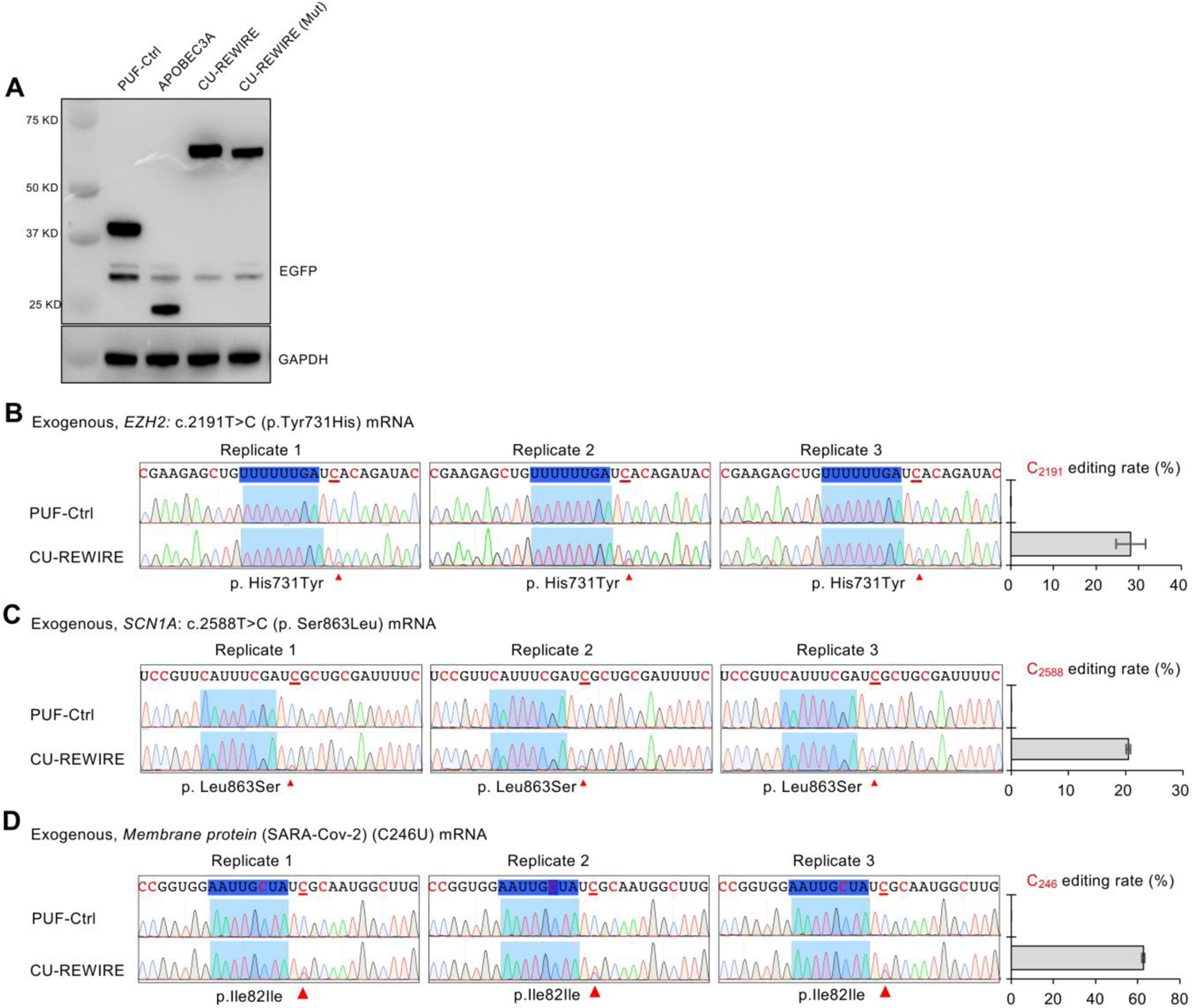
Protein expression level of CU-REWIREs and specific C-to-U base editing on exogenous pathogenetic mutant RNA. (A) Protein expression level of PUF-control, APOBEC3A-control and CU-REWIREs. Representative images of western blot were from the same samples in Figure 3B. The western blot was done in technical duplicate. (B) Example of using CU-REWIRE to correct disease-related mutation. The *EZH2* 2191T>C mutation is associated with Weaver syndrome disease, and we designed a CU-REWIRE to convert the 2191C back to U. The editing rates of all cytidines nearby the PUF binding site were measured by Sanger sequencing (Left), with the editing rate of the on-target of C2191 shown on the right. Values represent mean ± s.e.m. (n=3). (C) Engineering CU-REWIRE to correct *SCN1A* 2588T>C mutation which is associated with Weaver syndrome disease. The editing rates of all cytidines nearby the PUF binding site were measured by Sanger sequencing (Left), with the editing rate of the on-target of C2588 shown on the right. Values represent mean ± s.e.m. (n=3). (D) Engineering CU-REWIRE to editing M protein (SARS-Cov2). The editing rates of all cytidines nearby the PUF binding site were measured by Sanger sequencing (Left), with the editing rate of the on-target site C246 shown on the right. Values represent mean ± s.e.m. (n=3).

**Figure S6.**
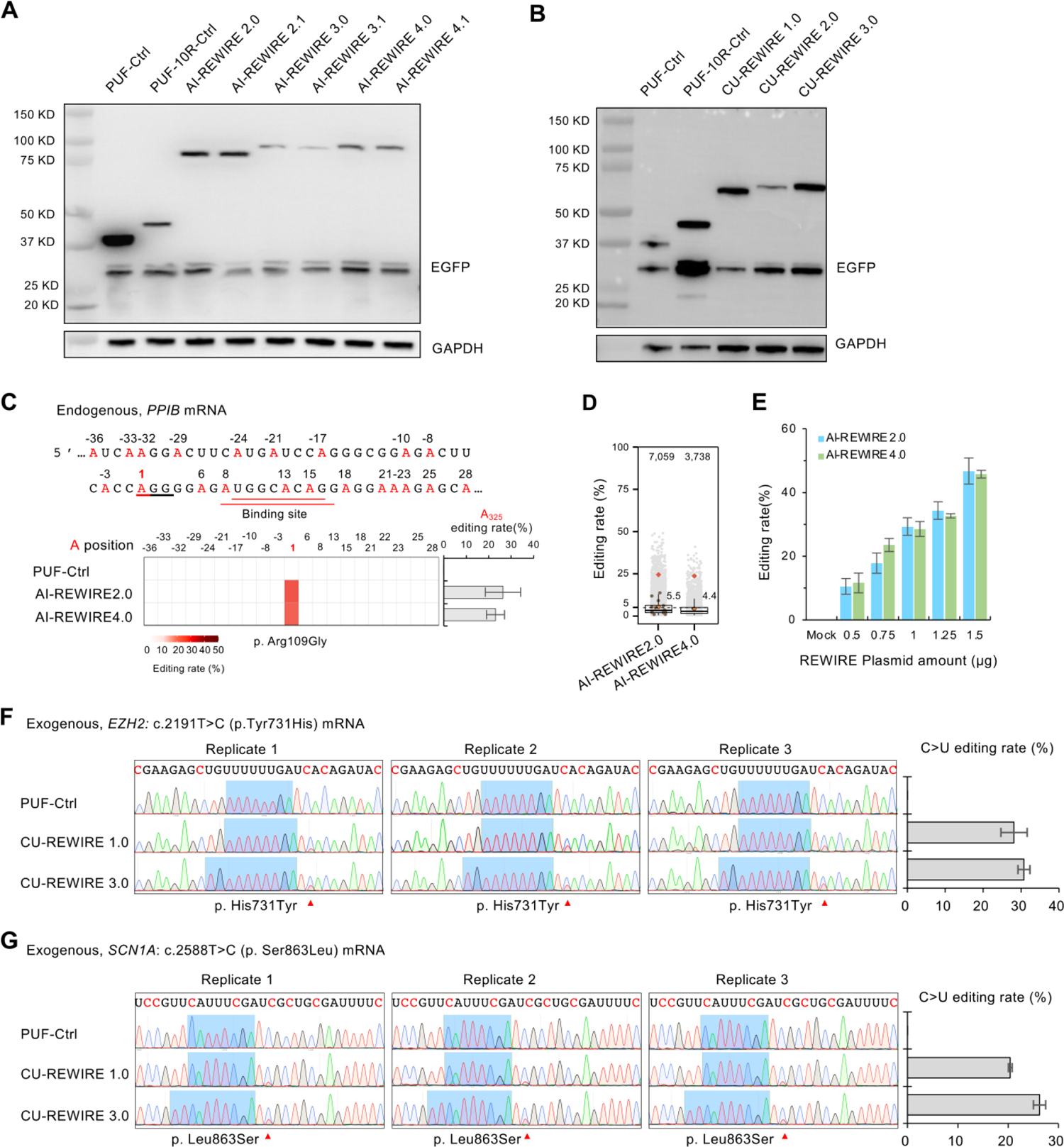
Protein expression level of upgraded REWIREs and assessment on RNA editing efficacy of original and upgraded REWIREs. (A) Protein expression levels of PUF-control and AI-REWIREs in HEK 293T cells. Representative images of western blot were the same samples in Figure 4B. The western blot was done in technical duplicate. (B) Western blot showed the protein expression levels of PUF-control and CU-REWIREs in HEK293T cells. Representative Western blot results used the same samples in Figure 4E. The western blot was done in technical duplicate. (C) Editing of *PPIB* mRNA by original and upgraded AI-REWIREs versions in HEK293T cell line. The PUF binding site is underlined in red, with the nearby adenosines marked in red. The editing rates of all adenosines nearby the PUF binding site were measured by RNA-seq (left), with the editing rate of the on-target site A325 shown on the right. Values represent mean ± s.e.m. (n=3). (D) Editing rates of transcriptome-wide A-to-I RNA editing in the samples treated with different AI-REWIREs (data from Figure S6C with minimum coverage cutoff at 100). The on-target *PPIB* site (325 A>G) is highlighted as red rhombuses, average editing rates of off-targets in each sample are indicated with orange dots and values are nearby; the 5% editing rate was indicated with a dashed line. The editing sites in endogenous genes with an identical PUF binding site sequence in close vicinity (+/- 20nt) were also marked in brown. (E) Bar plots showing the on-target RNA base editing rates of *PPIB* mRNA with different doses of original and upgraded AI-REWIREs expression vectors. The editing rates of the A325 site were tested by the Sanger sequencing. Values represent mean ± s.e.m. (n=3). (F) Correction of *EZH2* 2191T>C mutation associated with Weaver syndrome disease by CU-REWIREs. The PUF binding site is highlighted in blue, with the nearby cytidines marked in red. The editing rates of all cytidines nearby the PUF binding site were measured by Sanger sequencing (left), with the editing rate of on-target of C2191 shown in right. Values represent mean ± s.e.m. (n=3). (G) Correction of *SCN1A* 2588T>C mutation by CU-REWIREs. The editing rates of all cytidines nearby the PUF binding site were measured by Sanger sequencing (left), with the editing rate of on-target of C2588 shown in right. Values represent mean ± s.e.m. (n=3).

**Figure S7.**
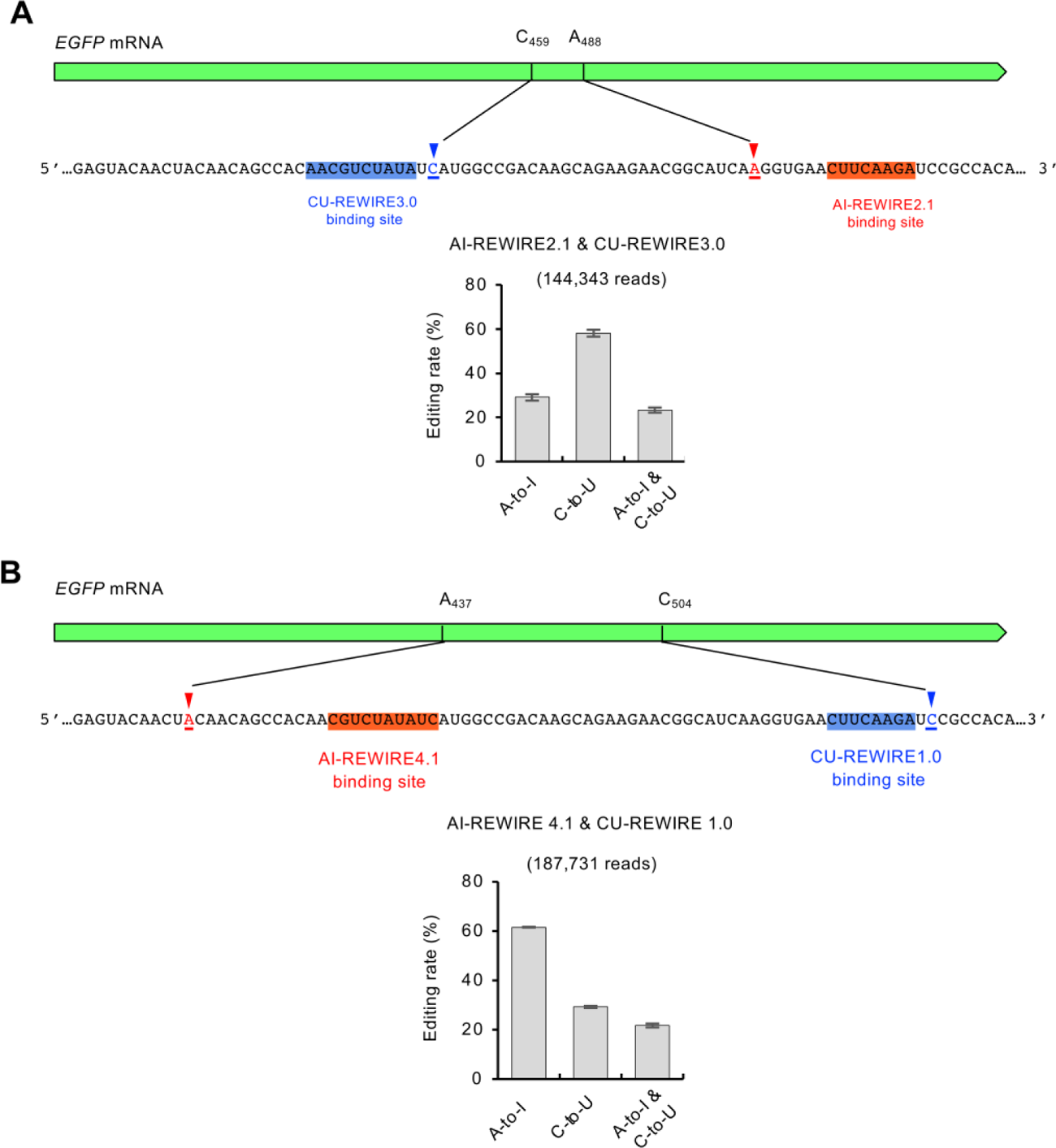
RNA base editing coordination mediated by AI&CU-REWIREs on the same transcript. (A) Schematic diagram of AI&CU-REWIREs achieved simultaneous base editing in the same transcript (top). Editing rates of the coordination with AI-REWIRE 2.1 and CU-REWIRE 3.0 measured by RNA-seq (bottom). (B) Schematic diagram of AI&CU-REWIREs targeted to different sites in the same EGFP transcript (top). Editing rates of the coordination with AI-REWIRE 4.1 and CU-REWIRE 1.0 measured by RNA-seq (bottom).

**Figure S8.**
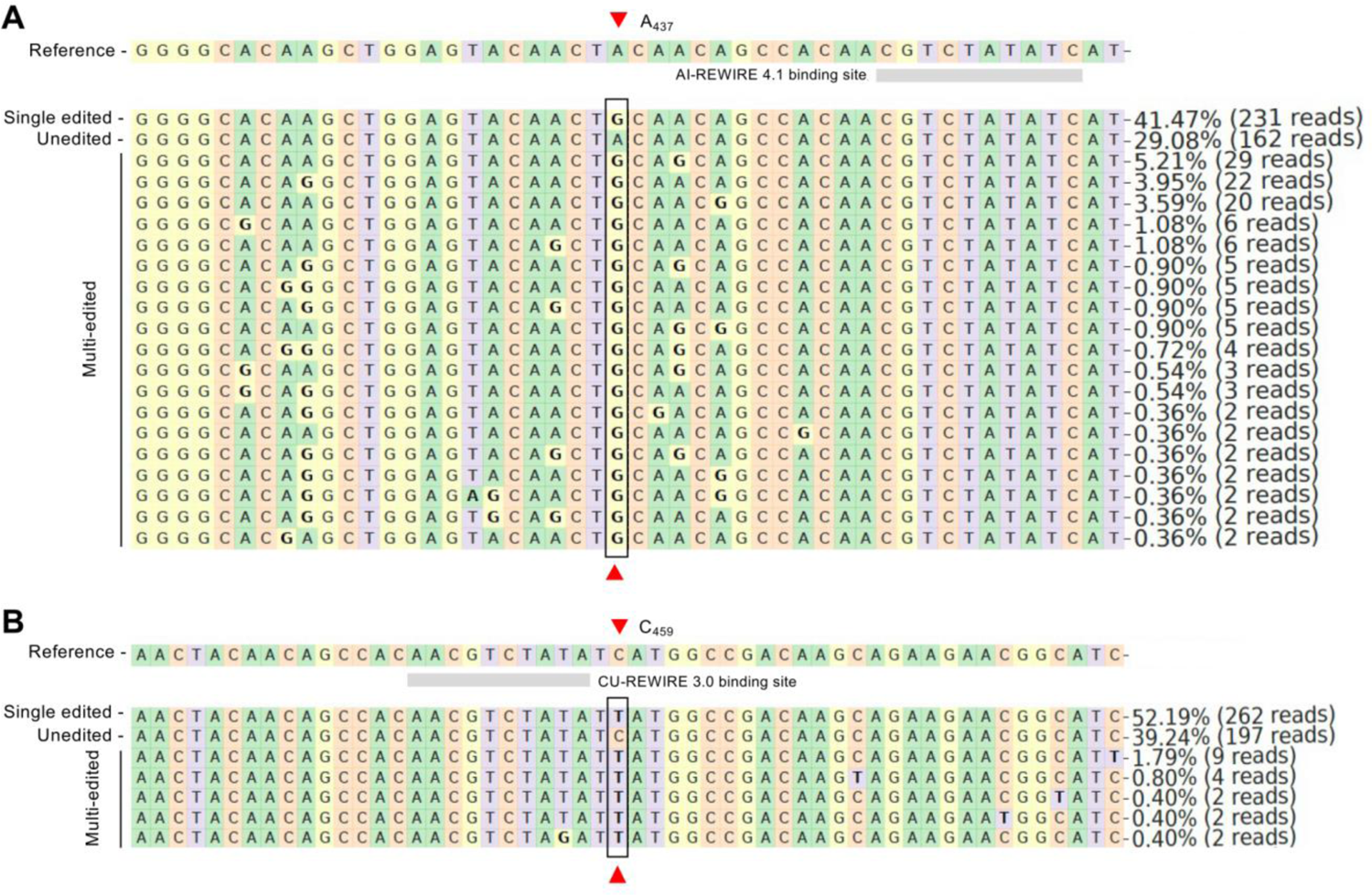
RNA editing allele table around target sites on EGFP mRNA by the AI-REWIRE 4.1 (A) and CU-REWIRE3.0 (B) in 293T cells. The target sites are indicated by red triangle. The PUF binding sites are underlined by grey tile. The percentile and number of reads of each sequence are listed. In this analysis, one representative data of three independent experiments was used; 200-nt window around targeted sites (100-nt upstream and downstream respectively) was used as reference; aligned reads must share ≥60% homology with reference sequence in Needleman-Wunsch alignment and have mapping quality ≥30; minimal depth cut to plot for one allele is 2. (A) AI-REWIRE4.1 replicate1; (B) CU-REIWIRE3.0 replicate1.

**Figure S9.**
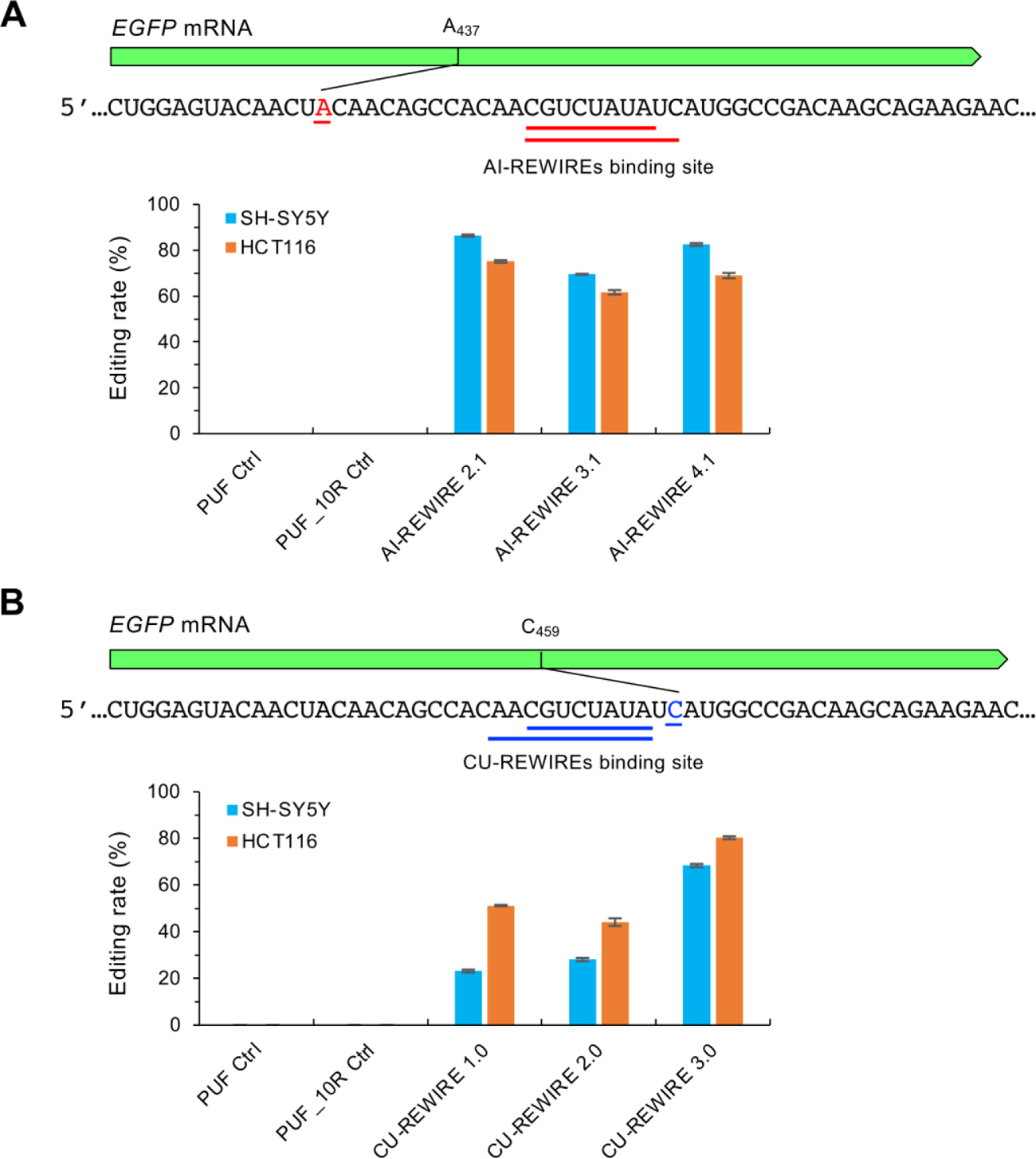
Application of REWIREs in other cancer cell lines. (A) Editing of EGFP mRNA by different AI-REWIRE versions in SH-SY5Y and HCT116 cell lines. The binding site of AI-REWIRE 2.1 in EGFP mRNA is underlined in black, AI-REWIRE 3.1/4.1 is underlined in red. The targeted cells were transfected with EGFP reporter and different versions of AI-REWIREs, and the total RNAs were extracted and purified 48 hours later. The editing rate of on-target site A437 was measured with Sanger sequencing (n=3). (B) Editing of EGFP mRNA by different versions of CU-REWIREs in SH-SY5Y and HCT116 cell line. The binding site of CU-REWIRE 1.0 in EGFP mRNA is underlined in black, CU-REWIRE 2.0/3.0 is underlined in blue. The targeted cells were transfected with EGFP reporter and different versions of CU-REWIREs, and the total RNAs were extracted and purified 48 hours later. The editing rate of on-target site C459 of EGFP mRNA was measured with Sanger sequencing (n=3).

**Figure S10.**
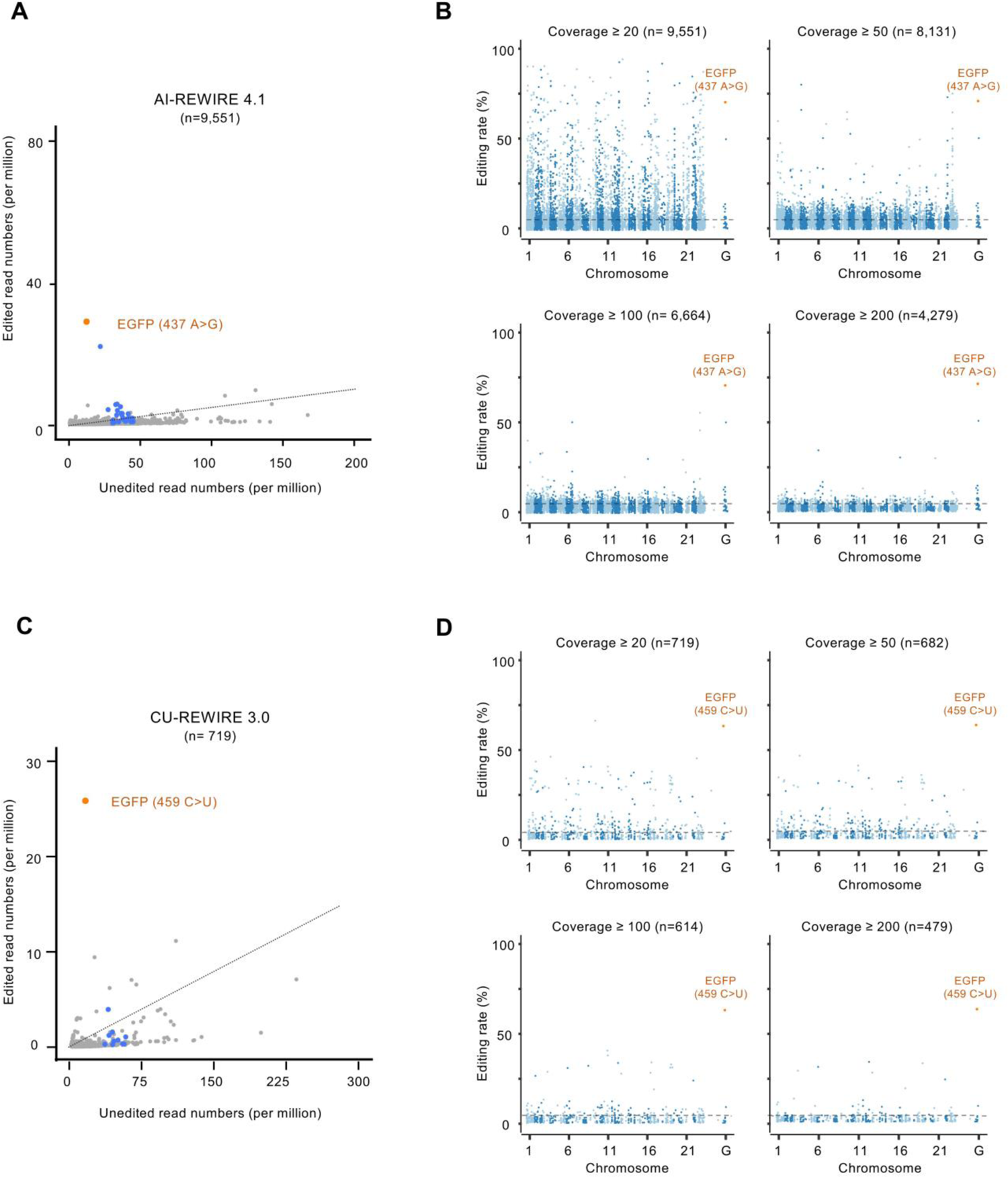
Comparative analysis of transcriptome-wide editing events with different coverage thresholds. (A) Scatter plots of transcriptome-wide A-to-I RNA editing sites in the samples treated with AI-REWIRE4.1. The total RNAs from each sample were sequenced using Illumina Hi-Seq X Ten. For each edited adenosine in human transcriptome, the numbers of unedited reads (x-axis) and edited reads (y-axis) were plotted. The experiments and analyses were carried out as described in Figure 1C with similar labels. (B) The off-target analysis across the entire transcriptome with different coverage thresholds. n, numbers of all adenosines detected. (C) Scatter plots of transcriptome-wide C-to-U RNA editing sites in the samples treated with CU-REWIRE3.0. The experiments and analyses were carried out as described in Figure 3C with similar labels. The off-target analysis across the entire transcriptome with different coverage thresholds. n, numbers of all cytosines detected.

## REFERENCES

1. Komor, A.C., Kim, Y.B., Packer, M.S., Zuris, J.A. and Liu, D.R. (2016) Programmable editing of a target base in genomic DNA without double-stranded DNA cleavage. Nature, 533, 420–424.

2. Wang, X., Li, J., Wang, Y., Yang, B., Wei, J., Wu, J., Wang, R., Huang, X., Chen, J. and Yang, L. (2018) Efficient base editing in methylated regions with a human APOBEC3A-Cas9 fusion. Nat Biotechnol, 36, 946–949.

3. Stafforst, T. and Schneider, M.F. (2012) An RNA-deaminase conjugate selectively repairs point mutations. Angew Chem Int Ed Engl, 51, 11166–11169.

4. Cox, D.B.T., Gootenberg, J.S., Abudayyeh, O.O., Franklin, B., Kellner, M.J., Joung, J. and Zhang, F. (2017) RNA editing with CRISPR-Cas13. Science, 358, 1019–1027.

5. Katrekar, D., Chen, G., Meluzzi, D., Ganesh, A., Worlikar, A., Shih, Y.R., Varghese, S. and Mali, P. (2019) In vivo RNA editing of point mutations via RNA-guided adenosine deaminases. Nat Methods, 16, 239–242.

6. Koblan, L.W., Erdos, M.R., Wilson, C., Cabral, W.A., Levy, J.M., Xiong, Z.M., Tavarez, U.L., Davison, L.M., Gete, Y.G., Mao, X. et al. (2021) In vivo base editing rescues Hutchinson-Gilford progeria syndrome in mice. Nature, 589, 608–614.

7. Erwei Zuo, Y.S., Wu Wei, Tanglong Yuan, Wenqin Ying, Hao Sun, Liyun Yuan4, Lars M. Steinmetz, Yixue Li, Hui Yang. (2019) Cytosine base editor generates substantial off-target single nucleotide variants in mouse embryos. science, 289–292.

8. Zhou, C., Sun, Y., Yan, R., Liu, Y., Zuo, E., Gu, C., Han, L., Wei, Y., Hu, X., Zeng, R. et al. (2019) Off-target RNA mutation induced by DNA base editing and its elimination by mutagenesis. Nature, 571, 275–278.

9. Grunewald, J., Zhou, R., Garcia, S.P., Iyer, S., Lareau, C.A., Aryee, M.J. and Joung, J.K. (2019) Transcriptome-wide off-target RNA editing induced by CRISPR-guided DNA base editors. Nature, 569, 433–437.

10. Kim, D., Lim, K., Kim, S.T., Yoon, S.H., Kim, K., Ryu, S.M. and Kim, J.S. (2017) Genome-wide target specificities of CRISPR RNA-guided programmable deaminases. Nat Biotechnol, 35, 475–480.

11. Montiel-Gonzalez, M.F., Vallecillo-Viejo, I., Yudowski, G.A. and Rosenthal, J.J. (2013) Correction of mutations within the cystic fibrosis transmembrane conductance regulator by site-directed RNA editing. Proc Natl Acad Sci U S A, 110, 18285–18290.

12. Fukuda, M., Umeno, H., Nose, K., Nishitarumizu, A., Noguchi, R. and Nakagawa, H. (2017) Construction of a guide-RNA for site-directed RNA mutagenesis utilising intracellular A-to-I RNA editing. Sci Rep, 7, 41478.

13. Wettengel, J., Reautschnig, P., Geisler, S., Kahle, P.J. and Stafforst, T. (2017) Harnessing human ADAR2 for RNA repair - Recoding a PINK1 mutation rescues mitophagy. Nucleic Acids Res, 45, 2797–2808.

14. Vogel, P., Moschref, M., Li, Q., Merkle, T., Selvasaravanan, K.D., Li, J.B. and Stafforst, T. (2018) Efficient and precise editing of endogenous transcripts with SNAP-tagged ADARs. Nat Methods, 15, 535–538.

15. Merkle, T., Merz, S., Reautschnig, P., Blaha, A., Li, Q., Vogel, P., Wettengel, J., Li, J.B. and Stafforst, T. (2019) Precise RNA editing by recruiting endogenous ADARs with antisense oligonucleotides. Nat Biotechnol, 37, 133–138.

16. Qu, L., Yi, Z., Zhu, S., Wang, C., Cao, Z., Zhou, Z., Yuan, P., Yu, Y., Tian, F., Liu, Z. et al. (2019) Programmable RNA editing by recruiting endogenous ADAR using engineered RNAs. Nat Biotechnol, 37, 1059–1069.

17. Abudayyeh, O.O., Gootenberg, J.S., Franklin, B., Koob, J., Kellner, M.J., Ladha, A., Joung, J., Kirchgatterer, P., Cox, D.B.T. and Zhang, F. (2019) A cytosine deaminase for programmable single-base RNA editing. Science, 365, 382–386.

18. Liu, Y., Mao, S., Huang, S., Li, Y., Chen, Y., Di, M., Huang, X., Lv, J., Wang, X., Ge, J. et al. (2020) REPAIRx, a specific yet highly efficient programmable A > I RNA base editor. EMBO J, 39, e104748.

19. Huang, X., Lv, J., Li, Y., Mao, S., Li, Z., Jing, Z., Sun, Y., Zhang, X., Shen, S., Wang, X. et al. (2020) Programmable C-to-U RNA editing using the human APOBEC3A deaminase. EMBO J, 39, e104741.

20. Rauch, S., He, E., Srienc, M., Zhou, H., Zhang, Z. and Dickinson, B.C. (2019) Programmable RNA-Guided RNA Effector Proteins Built from Human Parts. Cell, 178, 122–134 e112.

21. Mekler, V., Minakhin, L., Semenova, E., Kuznedelov, K. and Severinov, K. (2016) Kinetics of the CRISPR-Cas9 effector complex assembly and the role of 3’-terminal segment of guide RNA. Nucleic Acids Res, 44, 2837–2845.

22. Gough, V. and Gersbach, C.A. (2020) Immunity to Cas9 as an Obstacle to Persistent Genome Editing. Mol Ther, 28, 1389–1391.

23. Charlesworth, C.T., Deshpande, P.S., Dever, D.P., Camarena, J., Lemgart, V.T., Cromer, M.K., Vakulskas, C.A., Collingwood, M.A., Zhang, L., Bode, N.M. et al. (2019) Identification of preexisting adaptive immunity to Cas9 proteins in humans. Nat Med, 25, 249–254.

24. Wei, H. and Wang, Z. (2015) Engineering RNA-binding proteins with diverse activities. Wiley Interdiscip Rev RNA, 6, 597–613.

25. Cheong, C.G. and Hall, T.M. (2006) Engineering RNA sequence specificity of Pumilio repeats. Proc Natl Acad Sci U S A, 103, 13635–13639.

26. Dong, S., Wang, Y., Cassidy-Amstutz, C., Lu, G., Bigler, R., Jezyk, M.R., Li, C., Hall, T.M. and Wang, Z. (2011) Specific and modular binding code for cytosine recognition in Pumilio/FBF (PUF) RNA-binding domains. J Biol Chem, 286, 26732–26742.

27. Filipovska, A., Razif, M.F., Nygard, K.K. and Rackham, O. (2011) A universal code for RNA recognition by PUF proteins. Nat Chem Biol, 7, 425–427.

28. Ozawa, T., Natori, Y., Sato, M. and Umezawa, Y. (2007) Imaging dynamics of endogenous mitochondrial RNA in single living cells. Nat Methods, 4, 413–419.

29. Wang, Y., Cheong, C.G., Hall, T.M. and Wang, Z. (2009) Engineering splicing factors with designed specificities. Nat Methods, 6, 825–830.

30. Cooke, A., Prigge, A., Opperman, L. and Wickens, M. (2011) Targeted translational regulation using the PUF protein family scaffold. Proc Natl Acad Sci U S A, 108, 15870–15875.

31. Choudhury, R., Tsai, Y.S., Dominguez, D., Wang, Y. and Wang, Z. (2012) Engineering RNA endonucleases with customized sequence specificities. Nat Commun, 3, 1147.

32. Shinoda, K., Suda, A., Otonari, K., Futaki, S. and Imanishi, M. (2020) Programmable RNA methylation and demethylation using PUF RNA binding proteins. Chem Commun (Camb), 56, 1365–1368.

33. Adamala, K.P., Martin-Alarcon, D.A. and Boyden, E.S. (2016) Programmable RNA-binding protein composed of repeats of a single modular unit. Proc Natl Acad Sci U S A, 113, E2579–2588.

34. Salter, J.D., Bennett, R.P. and Smith, H.C. (2016) The APOBEC Protein Family: United by Structure, Divergent in Function. Trends Biochem Sci, 41, 578–594.

35. Zhao, Y.Y., Mao, M.W., Zhang, W.J., Wang, J., Li, H.T., Yang, Y., Wang, Z. and Wu, J.W. (2018) Expanding RNA binding specificity and affinity of engineered PUF domains. Nucleic Acids Res, 46, 4771–4782.

36. Jarmoskaite, I., Denny, S.K., Vaidyanathan, P.P., Becker, W.R., Andreasson, J.O.L., Layton, C.J., Kappel, K., Shivashankar, V., Sreenivasan, R., Das, R. et al. (2019) A Quantitative and Predictive Model for RNA Binding by Human Pumilio Proteins. Mol Cell, 74, 966–981 e918.

37. Wang, Y., Havel, J. and Beal, P.A. (2015) A Phenotypic Screen for Functional Mutants of Human Adenosine Deaminase Acting on RNA 1. ACS Chem Biol, 10, 2512–2519.

38. Kuttan, A. and Bass, B.L. (2012) Mechanistic insights into editing-site specificity of ADARs. Proc Natl Acad Sci U S A, 109, E3295–3304.

39. Li, H. and Durbin, R. (2009) Fast and accurate short read alignment with Burrows-Wheeler transform. Bioinformatics, 25, 1754–1760.

40. Li, H., Handsaker, B., Wysoker, A., Fennell, T., Ruan, J., Homer, N., Marth, G., Abecasis, G., Durbin, R. and Genome Project Data Processing, S. (2009) The Sequence Alignment/Map format and SAMtools. Bioinformatics, 25, 2078-2079.

41. Picardi, E., D’Erchia, A.M., Montalvo, A. and Pesole, G. (2015) Using REDItools to Detect RNA Editing Events in NGS Datasets. Curr Protoc Bioinformatics, 49, 12 12 11-12 12 15.

42. Dobin, A., Davis, C.A., Schlesinger, F., Drenkow, J., Zaleski, C., Jha, S., Batut, P., Chaisson, M. and Gingeras, T.R. (2013) STAR: ultrafast universal RNA-seq aligner. Bioinformatics, 29, 15–21.

43. Wagih, O. (2017) ggseqlogo: a versatile R package for drawing sequence logos. Bioinformatics, 33, 3645–3647.

44. Lehmann, K.A. and Bass, B.L. (2000) Double-stranded RNA adenosine deaminases ADAR1 and ADAR2 have overlapping specificities. Biochemistry, 39, 12875–12884.

45. Bohn, M.F., Shandilya, S.M.D., Silvas, T.V., Nalivaika, E.A., Kouno, T., Kelch, B.A., Ryder, S.P., Kurt-Yilmaz, N., Somasundaran, M. and Schiffer, C.A. (2015) The ssDNA Mutator APOBEC3A Is Regulated by Cooperative Dimerization. Structure, 23, 903–911.

46. Prasad, K.M., Xu, Y., Yang, Z., Acton, S.T. and French, B.A. (2011) Robust cardiomyocyte-specific gene expression following systemic injection of AAV: in vivo gene delivery follows a Poisson distribution. Gene Ther, 18, 43–52.

47. Zincarelli, C., Soltys, S., Rengo, G. and Rabinowitz, J.E. (2008) Analysis of AAV serotypes 1-9 mediated gene expression and tropism in mice after systemic injection. Mol Ther, 16, 1073–1080.

48. Gehrke, J.M., Cervantes, O., Clement, M.K., Wu, Y., Zeng, J., Bauer, D.E., Pinello, L. and Joung, J.K. (2018) An APOBEC3A-Cas9 base editor with minimized bystander and off-target activities. Nat Biotechnol, 36, 977–982.

49. Diroma, M.A., Ciaccia, L., Pesole, G. and Picardi, E. (2019) Elucidating the editome: bioinformatics approaches for RNA editing detection. Brief Bioinform, 20, 436–447.

50. Bahn, J.H., Lee, J.H., Li, G., Greer, C., Peng, G. and Xiao, X. (2012) Accurate identification of A-to-I RNA editing in human by transcriptome sequencing. Genome Res, 22, 142–150.

51. Bazak, L., Haviv, A., Barak, M., Jacob-Hirsch, J., Deng, P., Zhang, R., Isaacs, F.J., Rechavi, G., Li, J.B., Eisenberg, E. et al. (2014) A-to-I RNA editing occurs at over a hundred million genomic sites, located in a majority of human genes. Genome Res, 24, 365–376.

