## Supplementary Note 1. REWIREs plasmid protein sequences for "Programmable RNA base editing with a single gRNA-free enzyme"

REWIREs protein sequences used this study. Within REWIRE sequences, FLAG sequences are highlighted in yellow, PUF-8R or PUF-10R sequences are in green, ADAR or APOBEC3A sequences are in cyan.

**>PUF_8R (pCI-FLAG-PUF-8R(CGUCUAUA)):**

MDYKDDDDKGSRGRSRLLEDFRNNRYPNLQLREIAGHIMEFSQDQHGCRFIQLKLERATPAERQLVFNEILQAAYQLMVDVFGNYV
IQKFFEFGSLEQKLALAERIRGHVLSLALQMYGCRVIQKALEFIPSDQQNEMVRELDGHVLKCVKDQNGNHVVQKCIECVQPQSLQ
FIIDAFKGQVFALSTHPYGSYVIRRILEHCLPDQTLPILEELHQHTEQLVQDQYGNYVIQHVLEHGRPEDKSKIVAEIRGNVLVLS
QHKFASNVVEKCVTHASRTERAVLIDEVCTMNDGPHSALYTMMKDQYASYVVRKMIDVAEPGQRKIVMHKIRPHIATLRKYTYGKH
ILAKLEKYYMKNGVDLGSR*

**>PUF_10R (pCI-FLAG-PUF-10R(CGUCUAUAUC)):**

MDYKDDDDKGSRGRSRLLEDFRNNRYPNLQLREIAGHIMEFSQDQHGSYFIRLKLERATPAERQLVFNEILQAAYQLMVDVFGNYV
IQKFFEFGSLEQKLALAERIRGHVLSLALQMYGCRVIQKALEFIPSDQQNEMVRELDGQVFALSTHPYGNYVIQRILEHCLPDQTI
LEELHQHTEQLVQDQYGCRVIQHVLEHGRPEDKSKIVAEIRGNVLVLSQHKFANYVVQKCVTHASRTERAVLIDEVCTALYTMMKD
QYASYVVRKMIDVAEPGQRKIVMHKIRPHTEQLVQDQYGNYVIQHVLEHGRPEDKSKIVAEIRGNVLVLSQHKFASYVVEKCVTHA
SRTERAVLIDEVCTALYTMMKDQYASYVVRKMIDVAEPGQRKIVMHKIRPHIATLRKYTYGKHILAKLEKYYMKNGVDLGSR*

**PUF_10R* (pCI-FLAG- PUF-10R*(CGUCUAUAUC)):**

MDYKDDDDKGSRGRSRLLEDFRNNRYPNLQLREIAGHIMEFSQDQHGSYFIRLKLERATPAERQLVFNEILQAAYQLMVDVFGNYV
IQKFFEFGSLEQKLALAERIRGHVLSLALQMYGCRVIQKALEFIPSDQQNEMVRELDGQVFALSTHPYGNYVIQRILEHCLPDQTI
LEELHQHTEQLVQDQYGCRVIQHVLEHGRPEDKSKIVAEIRGNVLVLSQHKFANYVVQKCVTHASRTERAVLIDEVCTALYTMMKD
QYASYVVRKMIDVAEPGQRKIVMHKIRPHTEQLVQDQYGNYVIQHVLEHGRPEDKSKIVAEIRGNVLVLSQHKFASYVVEKCVTHA
SRTERAVLIDEVCTMNDGPHSALYTMMKDQYASYVVRKMIDVAEPGQRKIVMHKIRPHIATLRKYTYGKHILAKLEKYYMKNGVDL
GSR*

**>ADAR1 (pCI-FLAG-ADAR1$_{DD}$):**

MDYKDDDDKGSRVDEFKAERMGFTEVTPVTGASLRRTMLLLSRSPEAQPKTLPLTGSTFHDQIAMLSHRCFNTLTNSFQPSLLGRK
ILAAIIMKKDSEDMGVVVSLGTGNRCVKGDSLSLKGETVNDCHAEIISRRGFIRFLYSELMKYNSQTAKDSIFEPAKGGEKLQIKK
TVSFHLYISTAPCGDGALFDKSCSDRAMESTESRHYPVFENPKQGKLRTKVENGEGTIPVESSDIVPTWDGIRLGERLRTMSCSDK
ILRWNVLGLQGALLTHFLQPIYLKSVTLGYLFSQGHLTRAICCRVTRDGSAFEDGLRHPFIVNHPKVGRVSIYDSKRQSGKTKETS
VNWCLADGYDLEILDGTRGTVDGPRNELSRVSKKNIFLLFKKLCSFRYRRDLLRLSYGEAKKAARDYETAKNYFKKGLKDMGYGNW
ISKPQEEKNFYLCPV*

**>ADAR1* (pCI-FLAG-ADAR1*$_{DD}$):**

MDYKDDDDKGSRVDEFKAERMGFTEVTPVTGASLRRTMLLLSRSPEAQPKTLPLTGSTFHDQIAMLSHRCFNTLTNSFQPSLLGRK
ILAAIIMKKDSEDMGVVVSLGTGNRCVKGDSLSLKGETVNDCHAEIISRRGFIRFLYSELMKYNSQTAKDSIFEPAKGGEKLQIKK

TVSFHLYISTAPCGDGALFDKSCSDRAMESTESRHYPVFENPKQGKLRTKVENGQGTIPVESSDIVPTWDGIRLGERLRTMSCSDK
ILRWNVLGLQGALLTHFLQPIYLKSVTLGYLFSQGHLTRAICCRVTRDGSAFEDGLRHPFIVNHPKVGRVSIYDSKRQSGKTKETS
VNWCLADGYDLEILDGTRGTVDGPRNELSRVSKKNIFLLFKKLCSFRYRRDLLRLSYGEAKKAARDYETAKNYFKKGLKDMGYGNW
ISKPQEEKNFYLCPV*

**>ADAR2 (pCI-FLAG-ADAR2$_{DD}$):**

MDYKDDDDKGSRVDEFDQTPSRQPIPSEGLQLHLPQVLADAVSRLVLGKFGDLTDNFSSPHARRKVLAGVVMTTGTDVKDAKVISV
STGTKCINGEYMSDRGLALNDCHAEIISRRSLLRFLYTQLELYLNNKDDQKRSIFQKSERGGFRLKENVQFHLYISTSPCGDARIF
SPHEPILEGSRSYTQAGVQWCNHGSLQPRPPGLLSDPSTSTFQGAGTTEPADRHPNRKARGQLRTKIESGEGTIPVRSNASIQTWD
GVLQGERLLTMSCSDKIARWNVVGIQGSLLSIFVEPIYFSSIILGSLYHGDHLSRAMYQRISNIEDLPPLYTLNKPLLSGISNAEA
RQPGKAPNFSVNWTVGDSAIEVINATTGKDELGRASRLCKHALYCRWMRVHGKVPSHLLRSKITKPNVYHESKLAAKEYQAAKARL
FTAFIKAGLGAWVEKPTEQDQFSLTP*

**>ADAR2* (pCI-FLAG-ADAR2*$_{DD}$):**

MDYKDDDDKGSRVDEFDQTPSRQPIPSEGLQLHLPQVLADAVSRLVLGKFGDLTDNFSSPHARRKVLAGVVMTTGTDVKDAKVISV
STGTKCINGEYMSDRGLALNDCHAEIISRRSLLRFLYTQLELYLNNKDDQKRSIFQKSERGGFRLKENVQFHLYISTSPCGDARIF
SPHEPILEGSRSYTQAGVQWCNHGSLQPRPPGLLSDPSTSTFQGAGTTEPADRHPNRKARGQLRTKIESGQGTIPVRSNASIQTWD
GVLQGERLLTMSCSDKIARWNVVGIQGSLLSIFVEPIYFSSIILGSLYHGDHLSRAMYQRISNIEDLPPLYTLNKPLLSGISNAEA
RQPGKAPNFSVNWTVGDSAIEVINATTGKDELGRASRLCKHALYCRWMRVHGKVPSHLLRSKITKPNVYHESKLAAKEYQAAKARL
FTAFIKAGLGAWVEKPTEQDQFSLTP*

**>AI-REWIRE1.0 (pCI-FLAG-PUF-8R(CGUCUAUA)-ADAR1)**

MDYKDDDDKGSRGRSRLLEDFRNNRYPNLQLREIAGHIMEFSQDQHGSRFIQLKLERATPAERQLVFNEILQAAYQLMVDVFGNYV
IQKFFEFGSLEQKLALAERIRGHVLSLALQMYGCRVIQKALEFIPSDQQNEMVRELDGHVLKCVKDQNGNHVVQKCIECVQPQSLQ
FIIDAFKGQVFALSTHPYGSYVIRRILEHCLPDQTLPILEELHQHTEQLVQDQYGNYVIQHVLEHGRPEDKSKIVAEIRGNVLVLS
QHKFASNVVEKCVTHASRTERAVLIDEVCTMNDGPHSALYTMMKDQYASYVVRKMIDVAEPGQRKIVMHKIRPHIATLRKYTYGKH
ILAKLEKYYMKNGVDLGEFKAERMGFTEVTPVTGASLRRTMLLLSRSPEAQPKTLPLTGSTFHDQIAMLSHRCFNTLTNSFQPSLL
GRKILAAIIMKKDSEDMGVVVSLGTGNRCVKGDSLSLKGETVNDCHAEIISRRGFIRFLYSELMKYNSQTAKDSIFEPAKGGEKLQ
IKKTVSFHLYISTAPCGDGALFDKSCSDRAMESTESRHYPVFENPKQGKLRTKVENGEGTIPVESSDIVPTWDGIRLGERLRTMSC
SDKILRWNVLGLQGALLTHFLQPIYLKSVTLGYLFSQGHLTRAICCRVTRDGSAFEDGLRHPFIVNHPKVGRVSIYDSKRQSGKTK
ETSVNWCLADGYDLEILDGTRGTVDGPRNELSRVSKKNIFLLFKKLCSFRYRRDLLRLSYGEAKKAARDYETAKNYFKKGLKDMGY
GNWISKPQEEKNFYLCPV*.

**>AI-REWIRE2.0 (pCI-FLAG-PUF-8R(CGUCUAUA)-ADAR2)**

MDYKDDDDKGSRGRSRLLEDFRNNRYPNLQLREIAGHIMEFSQDQHGSRFIQLKLERATPAERQLVFNEILQAAYQLMVDVFGNYV
IQKFFEFGSLEQKLALAERIRGHVLSLALQMYGCRVIQKALEFIPSDQQNEMVRELDGHVLKCVKDQNGNHVVQKCIECVQPQSLQ

FIIDAFKGQVFALSTHPYGSYVIRRILEHCLPDQTLPILEELHQHTEQLVQDQYGNYVIQHVLEHGRPEDKSKIVAEIRGNVLVLS
QHKFASNVVEKCVTHASRTERAVLIDEVCTMNDGPHSALYTMMKDQYASYVVRKMIDVAEPGQRKIVMHKIRPHIATLRKYTYGKH
ILAKLEKYYMKNGVDLGEFDQTPSRQPIPSEGLQLHLPQVLADAVSRLVLGKFGDLTDNFSSPHARRKVLAGVVMTTGTDVKDAKV
ISVSTGTKCINGEYMSDRGLALNDCHAEIISRRSLLRFLYTQLELYLNNKDDQKRSIFQKSERGGFRLKENVQFHLYISTSPCGDA
RIFSPHEPILEGSRSYTQAGVQWCNHGSLQPRPPGLLSDPSTSTFQGAGTTEPADRHPNRKARGQLRTKIESGEGTIPVRSNASIQ
TWDGVLQGERLLTMSCSDKIARWNVVGIQGSLLSIFVEPIYFSSIILGSLYHGDHLSRAMYQRISNIEDLPPLYTLNKPLLSGISN
AEARQPGKAPNFSVNWTVGDSAIEVINATTGKDELGRASRLCKHALYCRWMRVHGKVPSHLLRSKITKPNVYHESKLAAKEYQAAK
ARLFTAFIKAGLGAWVEKPTEQDQFSLTP*

**>AI-REWIRE3.0 (pCI-FLAG-PUF-10R(CGUCUAUAUC)-ADAR2)**

MDYKDDDDKGSRGRSRLLEDFRNNRYPNLQLREIAGHIMEFSQDQHGSYFIRLKLERATPAERQLVFNEILQAAYQLMVDVFGNYV
IQKFFEFGSLEQKLALAERIRGHVLSLALQMYGCRVIQKALEFIPSDQQNEMVRELDGQVFALSTHPYGNYVIQRILEHCLPDQTI
LEELHQHTEQLVQDQYGCRVIQHVLEHGRPEDKSKIVAEIRGNVLVLSQHKFANYVVQKCVTHASRTERAVLIDEVCTALYTMMKD
QYASYVVRKMIDVAEPGQRKIVMHKIRPHTEQLVQDQYGNYVIQHVLEHGRPEDKSKIVAEIRGNVLVLSQHKFASYVVEKCVTHA
SRTERAVLIDEVCTALYTMMKDQYASYVVRKMIDVAEPGQRKIVMHKIRPHIATLRKYTYGKHILAKLEKYYMKNGVDLGEFDQTP
SRQPIPSEGLQLHLPQVLADAVSRLVLGKFGDLTDNFSSPHARRKVLAGVVMTTGTDVKDAKVISVSTGTKCINGEYMSDRGLALN
DCHAEIISRRSLLRFLYTQLELYLNNKDDQKRSIFQKSERGGFRLKENVQFHLYISTSPCGDARIFSPHEPILEGSRSYTQAGVQW
CNHGSLQPRPPGLLSDPSTSTFQGAGTTEPADRHPNRKARGQLRTKIESGEGTIPVRSNASIQTWDGVLQGERLLTMSCSDKIARW
NVVGIQGSLLSIFVEPIYFSSIILGSLYHGDHLSRAMYQRISNIEDLPPLYTLNKPLLSGISNAEARQPGKAPNFSVNWTVGDSAI
EVINATTGKDELGRASRLCKHALYCRWMRVHGKVPSHLLRSKITKPNVYHESKLAAKEYQAAKARLFTAFIKAGLGAWVEKPTEQD
QFSLTP*

**>AI-REWIRE4.0 (pCI-FLAG-PUF-10R*(CGUCUAUAUC)-ADAR2)**

MDYKDDDDKGSRGRSRLLEDFRNNRYPNLQLREIAGHIMEFSQDQHGSYFIRLKLERATPAERQLVFNEILQAAYQLMVDVFGNYV
IQKFFEFGSLEQKLALAERIRGHVLSLALQMYGCRVIQKALEFIPSDQQNEMVRELDGQVFALSTHPYGNYVIQRILEHCLPDQTI
LEELHQHTEQLVQDQYGCRVIQHVLEHGRPEDKSKIVAEIRGNVLVLSQHKFANYVVQKCVTHASRTERAVLIDEVCTALYTMMKD
QYASYVVRKMIDVAEPGQRKIVMHKIRPHTEQLVQDQYGNYVIQHVLEHGRPEDKSKIVAEIRGNVLVLSQHKFASYVVEKCVTHA
SRTERAVLIDEVCTMNDGPHSALYTMMKDQYASYVVRKMIDVAEPGQRKIVMHKIRPHIATLRKYTYGKHILAKLEKYYMKNGVDL
GEFDQTPSRQPIPSEGLQLHLPQVLADAVSRLVLGKFGDLTDNFSSPHARRKVLAGVVMTTGTDVKDAKVISVSTGTKCINGEYMS
DRGLALNDCHAEIISRRSLLRFLYTQLELYLNNKDDQKRSIFQKSERGGFRLKENVQFHLYISTSPCGDARIFSPHEPILEGSRSY
TQAGVQWCNHGSLQPRPPGLLSDPSTSTFQGAGTTEPADRHPNRKARGQLRTKIESGEGTIPVRSNASIQTWDGVLQGERLLTMSC
SDKIARWNVVGIQGSLLSIFVEPIYFSSIILGSLYHGDHLSRAMYQRISNIEDLPPLYTLNKPLLSGISNAEARQPGKAPNFSVNW
TVGDSAIEVINATTGKDELGRASRLCKHALYCRWMRVHGKVPSHLLRSKITKPNVYHESKLAAKEYQAAKARLFTAFIKAGLGAWV
EKPTEQDQFSLTP*

**>APOBEC3A (pCI-APOBEC3A-FLAG):**

MEASPASGPRHLMDPHIFTSNFNNGIGRHKTYLCYEVERLDNGTSVKMDQHRGFLHNQAKNLLCGFYGRHAELRFLDLVPSLQLDP

AQIYRVTWFISWSPCFSWGCAGEVRAFLQENTHVRLRIFAARIYDYDPLYKEALQMLRDAGAQVSIMTYDEFKHCWDTFVDHQGCP

FQPWDGLDEHSQALSGRLRAILQNQGNSGSETPGSRDYKDDDDK*

**>CU-REWIRE1.0 (pCI-APOBEC3A-XTEN-PUF-8R(CGUCUAUA)-FLAG)**

MEASPASGPRHLMDPHIFTSNFNNGIGRHKTYLCYEVERLDNGTSVKMDQHRGFLHNQAKNLLCGFYGRHAELRFLDLVPSLQLDP

AQIYRVTWFISWSPCFSWGCAGEVRAFLQENTHVRLRIFAARIYDYDPLYKEALQMLRDAGAQVSIMTYDEFKHCWDTFVDHQGCP

FQPWDGLDEHSQALSGRLRAILQNQGNSGSETPGTSESATPESGRSRLLEDFRNNRYPNLQLREIAGHIMEFSQDQHGSRFIQLKL

ERATPAERQLVFNEILQAAYQLMVDVFGNYVIQKFFEFGSLEQKLALAERIRGHVLSLALQMYGCRVIQKALEFIPSDQQNEMVRE

LDGHVLKCVKDQNGNHVVQKCIECVQPQSLQFIIDAFKGQVFALSTHPYGSYVIRRILEHCLPDQTLPILEELHQHTEQLVQDQYG

NYVIQHVLEHGRPEDKSKIVAEIRGNVLVLSQHKFASNVVEKCVTHASRTERAVLIDEVCTMNDGPHSALYTMMKDQYASYVVRKM

IDVAEPGQRKIVMHKIRPHIATLRKYTYGKHILAKLEKYYMKNGVDLGSRDYKDDDDK*

**>CU-REWIRE2.0 (pCI-APOBEC3A-XTEN-PUF-10(AACGUCUAUA)-FLAG)**

MEASPASGPRHLMDPHIFTSNFNNGIGRHKTYLCYEVERLDNGTSVKMDQHRGFLHNQAKNLLCGFYGRHAELRFLDLVPSLQLDP

AQIYRVTWFISWSPCFSWGCAGEVRAFLQENTHVRLRIFAARIYDYDPLYKEALQMLRDAGAQVSIMTYDEFKHCWDTFVDHQGCP

FQPWDGLDEHSQALSGRLRAILQNQGNSGSETPGTSESATPESGRSRLLEDFRNNRYPNLQLREIAGHIMEFSQDQHGCRFIQLKL

ERATPAERQLVFNEILQAAYQLMVDVFGNYVIQKFFEFGSLEQKLALAERIRGHVLSLALQMYGCRVIQKALEFIPSDQQNEMVRE

LDGQVFALSTHPYGNYVIQRILEHCLPDQTILEELHQHTEQLVQDQYGSYVIRHVLEHGRPEDKSKIVAEIRGNVLVLSQHKFANY

VVQKCVTHASRTERAVLIDEVCTALYTMMKDQYASYVVEKMIDVAEPGQRKIVMHKIRPHTEQLVQDQYGSYVIRHVLEHGRPEDK

SKIVAEIRGNVLVLSQHKFACRVVQKCVTHASRTERAVLIDEVCTALYTMMKDQYACRVVQKMIDVAEPGQRKIVMHKIRPHIATL

RKYTYGKHILAKLEKYYMKNGVDLGSRDYKDDDDK*

**>CU-REWIRE3.0 (pCI-APOBEC3A-XTEN-PUF-10*(AACGUCUAUA)-FLAG)**

MEASPASGPRHLMDPHIFTSNFNNGIGRHKTYLCYEVERLDNGTSVKMDQHRGFLHNQAKNLLCGFYGRHAELRFLDLVPSLQLDP

AQIYRVTWFISWSPCFSWGCAGEVRAFLQENTHVRLRIFAARIYDYDPLYKEALQMLRDAGAQVSIMTYDEFKHCWDTFVDHQGCP

FQPWDGLDEHSQALSGRLRAILQNQGNSGSETPGTSESATPESGRSRLLEDFRNNRYPNLQLREIAGHIMEFSQDQHGCRFIQLKL

ERATPAERQLVFNEILQAAYQLMVDVFGNYVIQKFFEFGSLEQKLALAERIRGHVLSLALQMYGCRVIQKALEFIPSDQQNEMVRE

LDGQVFALSTHPYGNYVIQRILEHCLPDQTILEELHQHTEQLVQDQYGSYVIRHVLEHGRPEDKSKIVAEIRGNVLVLSQHKFANY

VVQKCVTHASRTERAVLIDEVCTALYTMMKDQYASYVVEKMIDVAEPGQRKIVMHKIRPHTEQLVQDQYGSYVIRHVLEHGRPEDK

SKIVAEIRGNVLVLSQHKFACRVVQKCVTHASRTERAVLIDEVCTMNDGPHSALYTMMKDQYACRVVQKMIDVAEPGQRKIVMHKI

RPHIATLRKYTYGKHILAKLEKYYMKNGVDLGSRDYKDDDDK*

**Other PUF_8R and PUF_10R* sequence**

**> PUF_8R (GRIA2: GAUGCGAU)**

GRSRLLEDFRNNRYPNLQLREIAGHIMEFSQDQHGNYFIQLKLERATPAERQLVFNEILQAAYQLMVDVFGCRVIQKFFEFGSLEQ
KLALAERIRGHVLSLALQMYGSYVIEKALEFIPSDQQNEMVRELDGHVLKCVKDQNGNHVVQKCIECVQPQSLQFIIDAFKGQVFA
LSTHPYGSYVIERILEHCLPDQTLPILEELHQHTEQLVQDQYGNYVIQHVLEHGRPEDKSKIVAEIRGNVLVLSQHKFACRVVQKC
VTHASRTERAVLIDEVCTMNDGPHSALYTMMKDQYASYVVEKMIDVAEPGQRKIVMHKIRPHIATLRKYTYGKHILAKLEKYYMKN
GVDLG

**> PUF_8R (GRIA2: AUGCGAUA)**

GRSRLLEDFRNNRYPNLQLREIAGHIMEFSQDQHGCRFIQLKLERATPAERQLVFNEILQAAYQLMVDVFGNYVIQKFFEFGSLEQ
KLALAERIRGHVLSLALQMYGCRVIQKALEFIPSDQQNEMVRELDGHVLKCVKDQNGSYVVEKCIECVQPQSLQFIIDAFKGQVFA
LSTHPYGSYVIRRILEHCLPDQTLPILEELHQHTEQLVQDQYGSYVIEHVLEHGRPEDKSKIVAEIRGNVLVLSQHKFANYVVQKC
VTHASRTERAVLIDEVCTMNDGPHSALYTMMKDQYACRVVQKMIDVAEPGQRKIVMHKIRPHIATLRKYTYGKHILAKLEKYYMKN
GVDLG

**> PUF_8R (GRIA2: UGCGAUAU)**

GRSRLLEDFRNNRYPNLQLREIAGHIMEFSQDQHGNYFIQLKLERATPAERQLVFNEILQAAYQLMVDVFGCRVIQKFFEFGSLEQ
KLALAERIRGHVLSLALQMYGNYVIQKALEFIPSDQQNEMVRELDGHVLKCVKDQNGNHVVQKCIECVQPQSLQFIIDAFKGQVFA
LSTHPYGSYVIERILEHCLPDQTLPILEELHQHTEQLVQDQYGSYVIRHVLEHGRPEDKSKIVAEIRGNVLVLSQHKFASYVVEKC
VTHASRTERAVLIDEVCTMNDGPHSALYTMMKDQYANYVVQKMIDVAEPGQRKIVMHKIRPHIATLRKYTYGKHILAKLEKYYMKN
GVDLG

**> PUF_8R (GRIA2: GCGAUAUU)**

GRSRLLEDFRNNRYPNLQLREIAGHIMEFSQDQHGNYFIQLKLERATPAERQLVFNEILQAAYQLMVDVFGNYVIQKFFEFGSLEQ
KLALAERIRGHVLSLALQMYGCRVIQKALEFIPSDQQNEMVRELDGHVLKCVKDQNGNHVVQKCIECVQPQSLQFIIDAFKGQVFA
LSTHPYGCRVIQRILEHCLPDQTLPILEELHQHTEQLVQDQYGSYVIEHVLEHGRPEDKSKIVAEIRGNVLVLSQHKFASYVVRKC
VTHASRTERAVLIDEVCTMNDGPHSALYTMMKDQYASYVVEKMIDVAEPGQRKIVMHKIRPHIATLRKYTYGKHILAKLEKYYMKN
GVDLG

**> PUF_8R (GRIA2: CGAUAUUU)**

GRSRLLEDFRNNRYPNLQLREIAGHIMEFSQDQHGNYFIQLKLERATPAERQLVFNEILQAAYQLMVDVFGNYVIQKFFEFGSLEQ
KLALAERIRGHVLSLALQMYGNYVIQKALEFIPSDQQNEMVRELDGHVLKCVKDQNGNHVVQKCIECVQPQSLQFIIDAFKGQVFA
LSTHPYGNYVIQRILEHCLPDQTLPILEELHQHTEQLVQDQYGCRVIQHVLEHGRPEDKSKIVAEIRGNVLVLSQHKFASYVVEKC
VTHASRTERAVLIDEVCTMNDGPHSALYTMMKDQYASYVVRKMIDVAEPGQRKIVMHKIRPHIATLRKYTYGKHILAKLEKYYMKN
GVDLG

**> PUF_8R (GRIA2: GAUAUUUC)**

GRSRLLEDFRNNRYPNLQLREIAGHIMEFSQDQHGSYFIRLKLERATPAERQLVFNEILQAAYQLMVDVFGNYVIQKFFEFGSLEQ
KLALAERIRGHVLSLALQMYGNYVIQKALEFIPSDQQNEMVRELDGHVLKCVKDQNGNHVVQKCIECVQPQSLQFIIDAFKGQVFA
LSTHPYGCRVIQRILEHCLPDQTLPILEELHQHTEQLVQDQYGNYVIQHVLEHGRPEDKSKIVAEIRGNVLVLSQHKFACRVVQKC

VTHASRTERAVLIDEVCTMNDGPHSALYTMMKDQYASYVVEKMIDVAEPGQRKIVMHKIRPHIATLRKYTYGKHILAKLEKYYMKN

GVDLG

**> PUF_8R (GRIA2: AUAUUUCG)**

GRSRLLEDFRNNRYPNLQLREIAGHIMEFSQDQHGSYFIELKLERATPAERQLVFNEILQAAYQLMVDVFGSYVIRKFFEFGSLEQ

KLALAERIRGHVLSLALQMYGNYVIQKALEFIPSDQQNEMVRELDGHVLKCVKDQNGNHVVQKCIECVQPQSLQFIIDAFKGQVFA

LSTHPYGNYVIQRILEHCLPDQTLPILEELHQHTEQLVQDQYGCRVIQHVLEHGRPEDKSKIVAEIRGNVLVLSQHKFANYVVQKC

VTHASRTERAVLIDEVCTMNDGPHSALYTMMKDQYACRVVQKMIDVAEPGQRKIVMHKIRPHIATLRKYTYGKHILAKLEKYYMKN

GVDLG

**> PUF_8R (GRIA2: UAUUUCGC)**

GRSRLLEDFRNNRYPNLQLREIAGHIMEFSQDQHGSYFIRLKLERATPAERQLVFNEILQAAYQLMVDVFGSYVIEKFFEFGSLEQ

KLALAERIRGHVLSLALQMYGSYVIRKALEFIPSDQQNEMVRELDGHVLKCVKDQNGNHVVQKCIECVQPQSLQFIIDAFKGQVFA

LSTHPYGNYVIQRILEHCLPDQTLPILEELHQHTEQLVQDQYGNYVIQHVLEHGRPEDKSKIVAEIRGNVLVLSQHKFACRVVQKC

VTHASRTERAVLIDEVCTMNDGPHSALYTMMKDQYANYVVQKMIDVAEPGQRKIVMHKIRPHIATLRKYTYGKHILAKLEKYYMKN

GVDLG

**> PUF_8R (EGFP: CUUCAAGA)**

GRSRLLEDFRNNRYPNLQLREIAGHIMEFSQDQHGCRFIQLKLERATPAERQLVFNEILQAAYQLMVDVFGSYVIEKFFEFGSLEQ

KLALAERIRGHVLSLALQMYGCRVIQKALEFIPSDQQNEMVRELDGHVLKCVKDQNGNHVVQKCIECVQPQSLQFIIDAFKGQVFA

LSTHPYGSYVIRRILEHCLPDQTLPILEELHQHTEQLVQDQYGNYVIQHVLEHGRPEDKSKIVAEIRGNVLVLSQHKFANYVVQKC

VTHASRTERAVLIDEVCTMNDGPHSALYTMMKDQYASYVVRKMIDVAEPGQRKIVMHKIRPHIATLRKYTYGKHILAKLEKYYMKN

GVDLG

**> PUF_8R (TP53: UCUGGCCC)**

GRSRLLEDFRNNRYPNLQLREIAGHIMEFSQDQHGSYFIRLKLERATPAERQLVFNEILQAAYQLMVDVFGSYVIRKFFEFGSLEQ

KLALAERIRGHVLSLALQMYGSYVIRKALEFIPSDQQNEMVRELDGHVLKCVKDQNGSYVVEKCIECVQPQSLQFIIDAFKGQVFA

LSTHPYGSYVIERILEHCLPDQTLPILEELHQHTEQLVQDQYGNYVIQHVLEHGRPEDKSKIVAEIRGNVLVLSQHKFASYVVRKC

VTHASRTERAVLIDEVCTMNDGPHSALYTMMKDQYANYVVQKMIDVAEPGQRKIVMHKIRPHIATLRKYTYGKHILAKLEKYYMKN

GVDLG

**> PUF_8R (TP53: GGCGCUGC)**

GRSRLLEDFRNNRYPNLQLREIAGHIMEFSQDQHGSYFIRLKLERATPAERQLVFNEILQAAYQLMVDVFGSYVIEKFFEFGSLEQ

KLALAERIRGHVLSLALQMYGNYVIQKALEFIPSDQQNEMVRELDGHVLKCVKDQNGNHVVQKCIECVQPQSLQFIIDAFKGQVFA

LSTHPYGSYVIERILEHCLPDQTLPILEELHQHTEQLVQDQYGSYVIRHVLEHGRPEDKSKIVAEIRGNVLVLSQHKFASYVVEKC

VTHASRTERAVLIDEVCTMNDGPHSALYTMMKDQYASYVVEKMIDVAEPGQRKIVMHKIRPHIATLRKYTYGKHILAKLEKYYMKN

GVDLG

**> PUF_8R (PPIB: TCAAGGTG)**

GRSRLLEDFRNNRYPNLQLREIAGHIMEFSQDQHGSYFIELKLERATPAERQLVFNEILQAAYQLMVDVFGNYVIQKFFEFGSLEQ
KLALAERIRGHVLSLALQMYGSYVIEKALEFIPSDQQNEMVRELDGHVLKCVKDQNGSYVVEKCIECVQPQSLQFIIDAFKGQVFA
LSTHPYGCRVIQRILEHCLPDQTLPILEELHQHTEQLVQDQYGCRVIQHVLEHGRPEDKSKIVAEIRGNVLVLSQHKFASYVVRKC
VTHASRTERAVLIDEVCTMNDGPHSALYTMMKDQYANYVVQKMIDVAEPGQRKIVMHKIRPHIATLRKYTYGKHILAKLEKYYMKN
GVDLG

**> PUF_8R (PPIB: TGGCACAG)**

GRSRLLEDFRNNRYPNLQLREIAGHIMEFSQDQHGSYFIELKLERATPAERQLVFNEILQAAYQLMVDVFGCRVIQKFFEFGSLEQ
KLALAERIRGHVLSLALQMYGSYVIRKALEFIPSDQQNEMVRELDGHVLKCVKDQNGCRVVQKCIECVQPQSLQFIIDAFKGQVFA
LSTHPYGSYVIRRILEHCLPDQTLPILEELHQHTEQLVQDQYGSYVIEHVLEHGRPEDKSKIVAEIRGNVLVLSQHKFASYVVEKC
VTHASRTERAVLIDEVCTMNDGPHSALYTMMKDQYANYVVQKMIDVAEPGQRKIVMHKIRPHIATLRKYTYGKHILAKLEKYYMKN
GVDLG

**> PUF_8R (BMPR2: CGTCTTGC)**

GRSRLLEDFRNNRYPNLQLREIAGHIMEFSQDQHGSYFIRLKLERATPAERQLVFNEILQAAYQLMVDVFGSYVIEKFFEFGSLEQ
KLALAERIRGHVLSLALQMYGNYVIQKALEFIPSDQQNEMVRELDGHVLKCVKDQNGNHVVQKCIECVQPQSLQFIIDAFKGQVFA
LSTHPYGSYVIRRILEHCLPDQTLPILEELHQHTEQLVQDQYGNYVIQHVLEHGRPEDKSKIVAEIRGNVLVLSQHKFASYVVEKC
VTHASRTERAVLIDEVCTMNDGPHSALYTMMKDQYASYVVRKMIDVAEPGQRKIVMHKIRPHIATLRKYTYGKHILAKLEKYYMKN
GVDLG

**> PUF_8R (DMD: TTACAAGA)**

GRSRLLEDFRNNRYPNLQLREIAGHIMEFSQDQHGCRFIQLKLERATPAERQLVFNEILQAAYQLMVDVFGSYVIEKFFEFGSLEQ
KLALAERIRGHVLSLALQMYGCRVIQKALEFIPSDQQNEMVRELDGHVLKCVKDQNGNHVVQKCIECVQPQSLQFIIDAFKGQVFA
LSTHPYGSYVIRRILEHCLPDQTLPILEELHQHTEQLVQDQYGCRVIQHVLEHGRPEDKSKIVAEIRGNVLVLSQHKFANYVVQKC
VTHASRTERAVLIDEVCTMNDGPHSALYTMMKDQYANYVVQKMIDVAEPGQRKIVMHKIRPHIATLRKYTYGKHILAKLEKYYMKN
GVDLG

**> PUF_8R (DMD: TTACTGAA)**

GRSRLLEDFRNNRYPNLQLREIAGHIMEFSQDQHGCRFIQLKLERATPAERQLVFNEILQAAYQLMVDVFGCRVIQKFFEFGSLEQ
KLALAERIRGHVLSLALQMYGSYVIEKALEFIPSDQQNEMVRELDGHVLKCVKDQNGNHVVQKCIECVQPQSLQFIIDAFKGQVFA
LSTHPYGSYVIRRILEHCLPDQTLPILEELHQHTEQLVQDQYGCRVIQHVLEHGRPEDKSKIVAEIRGNVLVLSQHKFANYVVQKC
VTHASRTERAVLIDEVCTMNDGPHSALYTMMKDQYANYVVQKMIDVAEPGQRKIVMHKIRPHIATLRKYTYGKHILAKLEKYYMKN
GVDLG

**> PUF_8R (COL3A1: TGGTTCCC)**

GRSRLLEDFRNNRYPNLQLREIAGHIMEFSQDQHGSYFIRLKLERATPAERQLVFNEILQAAYQLMVDVFGSYVIRKFFEFGSLEQ
KLALAERIRGHVLSLALQMYGSYVIRKALEFIPSDQQNEMVRELDGHVLKCVKDQNGNHVVQKCIECVQPQSLQFIIDAFKGQVFA
LSTHPYGNYVIQRILEHCLPDQTLPILEELHQHTEQLVQDQYGSYVIEHVLEHGRPEDKSKIVAEIRGNVLVLSQHKFASYVVEKC

VTHASRTERAVLIDEVCTMNDGPHSALYTMMKDQYANYVVQKMIDVAEPGQRKIVMHKIRPHIATLRKYTYGKHILAKLEKYYMKN

GVDLG

**> PUF_8R (COL3A1: GGACCTCC)**

GRSRLLEDFRNNRYPNLQLREIAGHIMEFSQDQHGSYFIRLKLERATPAERQLVFNEILQAAYQLMVDVFGSYVIRKFFEFGSLEQ

KLALAERIRGHVLSLALQMYGNYVIQKALEFIPSDQQNEMVRELDGHVLKCVKDQNGNHVVQKCIECVQPQSLQFIIDAFKGQVFA

LSTHPYGSYVIRRILEHCLPDQTLPILEELHQHTEQLVQDQYGCRVIQHVLEHGRPEDKSKIVAEIRGNVLVLSQHKFASYVVEKC

VTHASRTERAVLIDEVCTMNDGPHSALYTMMKDQYASYVVEKMIDVAEPGQRKIVMHKIRPHIATLRKYTYGKHILAKLEKYYMKN

GVDLG

**> PUF_8R (HBB: TGAACGTG)**

GRSRLLEDFRNNRYPNLQLREIAGHIMEFSQDQHGSYFIELKLERATPAERQLVFNEILQAAYQLMVDVFGNYVIQKFFEFGSLEQ

KLALAERIRGHVLSLALQMYGSYVIEKALEFIPSDQQNEMVRELDGHVLKCVKDQNGNHVVQKCIECVQPQSLQFIIDAFKGQVFA

LSTHPYGCRVIQRILEHCLPDQTLPILEELHQHTEQLVQDQYGCRVIQHVLEHGRPEDKSKIVAEIRGNVLVLSQHKFASYVVEKC

VTHASRTERAVLIDEVCTMNDGPHSALYTMMKDQYANYVVQKMIDVAEPGQRKIVMHKIRPHIATLRKYTYGKHILAKLEKYYMKN

GVDLG

**> PUF_8R (HBB: GTTCTTTG)**

GRSRLLEDFRNNRYPNLQLREIAGHIMEFSQDQHGSYFIELKLERATPAERQLVFNEILQAAYQLMVDVFGNYVIQKFFEFGSLEQ

KLALAERIRGHVLSLALQMYGNYVIQKALEFIPSDQQNEMVRELDGHVLKCVKDQNGNHVVQKCIECVQPQSLQFIIDAFKGQVFA

LSTHPYGSYVIRRILEHCLPDQTLPILEELHQHTEQLVQDQYGNYVIQHVLEHGRPEDKSKIVAEIRGNVLVLSQHKFANYVVQKC

VTHASRTERAVLIDEVCTMNDGPHSALYTMMKDQYASYVVEKMIDVAEPGQRKIVMHKIRPHIATLRKYTYGKHILAKLEKYYMKN

GVDLG

**> PUF_8R (EGFP: AACUUCAA)**

GRSRLLEDFRNNRYPNLQLREIAGHIMEFSQDQHGCRFIQLKLERATPAERQLVFNEILQAAYQLMVDVFGCRVIQKFFEFGSLEQ

KLALAERIRGHVLSLALQMYGSYVIRKALEFIPSDQQNEMVRELDGHVLKCVKDQNGNHVVQKCIECVQPQSLQFIIDAFKGQVFA

LSTHPYGNYVIQRILEHCLPDQTLPILEELHQHTEQLVQDQYGSYVIRHVLEHGRPEDKSKIVAEIRGNVLVLSQHKFACRVVQKC

VTHASRTERAVLIDEVCTMNDGPHSALYTMMKDQYACRVVQKMIDVAEPGQRKIVMHKIRPHIATLRKYTYGKHILAKLEKYYMKN

GVDLG

**> PUF_8R (EGFP: ACUUCAAG)**

GRSRLLEDFRNNRYPNLQLREIAGHIMEFSQDQHGSYFIELKLERATPAERQLVFNEILQAAYQLMVDVFGCRVIQKFFEFGSLEQ

KLALAERIRGHVLSLALQMYGCRVIQKALEFIPSDQQNEMVRELDGHVLKCVKDQNGNHVVQKCIECVQPQSLQFIIDAFKGQVFA

LSTHPYGNYVIQRILEHCLPDQTLPILEELHQHTEQLVQDQYGNYVIQHVLEHGRPEDKSKIVAEIRGNVLVLSQHKFASYVVRKC

VTHASRTERAVLIDEVCTMNDGPHSALYTMMKDQYACRVVQKMIDVAEPGQRKIVMHKIRPHIATLRKYTYGKHILAKLEKYYMKN

GVDLG

**> PUF_8R (EGFP: UUCAAGAU)**

GRSRLLEDFRNNRYPNLQLREIAGHIMEFSQDQHGNYFIQLKLERATPAERQLVFNEILQAAYQLMVDVFGCRVIQKFFEFGSLEQ
KLALAERIRGHVLSLALQMYGSYVIEKALEFIPSDQQNEMVRELDGHVLKCVKDQNGCRVVQKCIECVQPQSLQFIIDAFKGQVFA
LSTHPYGCRVIQRILEHCLPDQTLPILEELHQHTEQLVQDQYGSYVIRHVLEHGRPEDKSKIVAEIRGNVLVLSQHKFANYVVQKC
VTHASRTERAVLIDEVCTMNDGPHSALYTMMKDQYANYVVQKMIDVAEPGQRKIVMHKIRPHIATLRKYTYGKHILAKLEKYYMKN
GVDLG

**> PUF_8R (EGFP: UCAAGAUC)**

GRSRLLEDFRNNRYPNLQLREIAGHIMEFSQDQHGSYFIRLKLERATPAERQLVFNEILQAAYQLMVDVFGNYVIQKFFEFGSLEQ
KLALAERIRGHVLSLALQMYGCRVIQKALEFIPSDQQNEMVRELDGHVLKCVKDQNGSYVVEKCIECVQPQSLQFIIDAFKGQVFA
LSTHPYGCRVIQRILEHCLPDQTLPILEELHQHTEQLVQDQYGCRVIQHVLEHGRPEDKSKIVAEIRGNVLVLSQHKFASYVVRKC
VTHASRTERAVLIDEVCTMNDGPHSALYTMMKDQYANYVVQKMIDVAEPGQRKIVMHKIRPHIATLRKYTYGKHILAKLEKYYMKN
GVDLG

**> PUF_8R (EZH2: UUUUUUGA)**

GRSRLLEDFRNNRYPNLQLREIAGHIMEFSQDQHGCRFIQLKLERATPAERQLVFNEILQAAYQLMVDVFGSYVIEKFFEFGSLEQ
KLALAERIRGHVLSLALQMYGNYVIQKALEFIPSDQQNEMVRELDGHVLKCVKDQNGNHVVQKCIECVQPQSLQFIIDAFKGQVFA
LSTHPYGNYVIQRILEHCLPDQTLPILEELHQHTEQLVQDQYGNYVIQHVLEHGRPEDKSKIVAEIRGNVLVLSQHKFANYVVQKC
VTHASRTERAVLIDEVCTMNDGPHSALYTMMKDQYANYVVQKMIDVAEPGQRKIVMHKIRPHIATLRKYTYGKHILAKLEKYYMKN
GVDLG

**> PUF_8R (SCN1A: CAUUUCGA)**

GRSRLLEDFRNNRYPNLQLREIAGHIMEFSQDQHGCRFIQLKLERATPAERQLVFNEILQAAYQLMVDVFGSYVIEKFFEFGSLEQ
KLALAERIRGHVLSLALQMYGSYVIRKALEFIPSDQQNEMVRELDGHVLKCVKDQNGNHVVQKCIECVQPQSLQFIIDAFKGQVFA
LSTHPYGNYVIQRILEHCLPDQTLPILEELHQHTEQLVQDQYGNYVIQHVLEHGRPEDKSKIVAEIRGNVLVLSQHKFACRVVQKC
VTHASRTERAVLIDEVCTMNDGPHSALYTMMKDQYASYVVRKMIDVAEPGQRKIVMHKIRPHIATLRKYTYGKHILAKLEKYYMKN
GVDLG

**> PUF_8R (CTNNB1: AUUCGAAA)**

GRSRLLEDFRNNRYPNLQLREIAGHIMEFSQDQHGCRFIQLKLERATPAERQLVFNEILQAAYQLMVDVFGCRVIQKFFEFGSLEQ
KLALAERIRGHVLSLALQMYGCRVIQKALEFIPSDQQNEMVRELDGHVLKCVKDQNGSYVVEKCIECVQPQSLQFIIDAFKGQVFA
LSTHPYGSYVIERILEHCLPDQTLPILEELHQHTEQLVQDQYGNYVIQHVLEHGRPEDKSKIVAEIRGNVLVLSQHKFANYVVQKC
VTHASRTERAVLIDEVCTMNDGPHSALYTMMKDQYACRVVQKMIDVAEPGQRKIVMHKIRPHIATLRKYTYGKHILAKLEKYYMKN
GVDLG

**> PUF_8R (SARS-COV2: AAUUGCUA)**

GRSRLLEDFRNNRYPNLQLREIAGHIMEFSQDQHGCRFIQLKLERATPAERQLVFNEILQAAYQLMVDVFGNYVIQKFFEFGSLEQ
KLALAERIRGHVLSLALQMYGSYVIRKALEFIPSDQQNEMVRELDGHVLKCVKDQNGSYVVEKCIECVQPQSLQFIIDAFKGQVFA
LSTHPYGNYVIQRILEHCLPDQTLPILEELHQHTEQLVQDQYGNYVIQHVLEHGRPEDKSKIVAEIRGNVLVLSQHKFACRVVQKC

VTHASRTERAVLIDEVCTMNDGPHSALYTMMKDQYACRVVQKMIDVAEPGQRKIVMHKIRPHIATLRKYTYGKHILAKLEKYYMKN
GVDLG

**> PUF_10R* (EGFP: AACGUCUAUA)**

GRSRLLEDFRNNRYPNLQLREIAGHIMEFSQDQHGCRFIQLKLERATPAERQLVFNEILQAAYQLMVDVFGNYVIQKFFEFGSLEQ
KLALAERIRGHVLSLALQMYGCRVIQKALEFIPSDQQNEMVRELDGQVFALSTHPYGNYVIQRILEHCLPDQTILEELHQHTEQLV
QDQYGNYVIQHVLEHGRPEDKSKIVAEIRGNVLVLSQHKFANYVVQKCVTHASRTERAVLIDEVCTALYTMMKDQYASYVVEKMID
VAEPGQRKIVMHKIRPHTEQLVQDQYGSYVIRHVLEHGRPEDKSKIVAEIRGNVLVLSQHKFACRVVQKCVTHASRTERAVLIDEV
CTMNDGPHSALYTMMKDQYACRVVQKMIDVAEPGQRKIVMHKIRPHIATLRKYTYGKHILAKLEKYYMKNGVDLG

**> PUF_10R* (PPIB: AUGGCACAGG)**

GRSRLLEDFRNNRYPNLQLREIAGHIMEFSQDQHGSYFIELKLERATPAERQLVFNEILQAAYQLMVDVFGSYVIEKFFEFGSLEQ
KLALAERIRGHVLSLALQMYGCRVIQKALEFIPSDQQNEMVRELDGQVFALSTHPYGSYVIRRILEHCLPDQTILEELHQHTEQLV
QDQYGCRVIQHVLEHGRPEDKSKIVAEIRGNVLVLSQHKFASYVVRKCVTHASRTERAVLIDEVCTALYTMMKDQYASYVVEKMID
VAEPGQRKIVMHKIRPHTEQLVQDQYGSYVIEHVLEHGRPEDKSKIVAEIRGNVLVLSQHKFANYVVQKCVTHASRTERAVLIDEV
CTMNDGPHSALYTMMKDQYACRVVQKMIDVAEPGQRKIVMHKIRPHIATLRKYTYGKHILAKLEKYYMKNGVDLG

**> PUF_10R* (EZH2: UGUUUUUUGA)**

GRSRLLEDFRNNRYPNLQLREIAGHIMEFSQDQHGCRFIQLKLERATPAERQLVFNEILQAAYQLMVDVFGSYVIEKFFEFGSLEQ
KLALAERIRGHVLSLALQMYGNYVIQKALEFIPSDQQNEMVRELDGQVFALSTHPYGNYVIQRILEHCLPDQTILEELHQHTEQLV
QDQYGNYVIQHVLEHGRPEDKSKIVAEIRGNVLVLSQHKFANYVVQKCVTHASRTERAVLIDEVCTALYTMMKDQYANYVVQKMID
VAEPGQRKIVMHKIRPHTEQLVQDQYGNYVIQHVLEHGRPEDKSKIVAEIRGNVLVLSQHKFASYVVEKCVTHASRTERAVLIDEV
CTMNDGPHSALYTMMKDQYANYVVQKMIDVAEPGQRKIVMHKIRPHIATLRKYTYGKHILAKLEKYYMKNGVDLG

**> PUF_10R* (SCN1A: UUCAUUUCGA)**

GRSRLLEDFRNNRYPNLQLREIAGHIMEFSQDQHGCRFIQLKLERATPAERQLVFNEILQAAYQLMVDVFGSYVIEKFFEFGSLEQ
KLALAERIRGHVLSLALQMYGSYVIRKALEFIPSDQQNEMVRELDGQVFALSTHPYGNYVIQRILEHCLPDQTILEELHQHTEQLV
QDQYGNYVIQHVLEHGRPEDKSKIVAEIRGNVLVLSQHKFANYVVQKCVTHASRTERAVLIDEVCTALYTMMKDQYACRVVQKMID
VAEPGQRKIVMHKIRPHTEQLVQDQYGSYVIRHVLEHGRPEDKSKIVAEIRGNVLVLSQHKFANYVVQKCVTHASRTERAVLIDEV
CTMNDGPHSALYTMMKDQYANYVVQKMIDVAEPGQRKIVMHKIRPHIATLRKYTYGKHILAKLEKYYMKNGVDLG.
