## Supplementary material for "Programmable RNA base editing with a single gRNA-free enzyme": Table S1

**Table S1 Primes, minigenes and plasmids are used in this study.**

| Gene name | Primer name | Primer sequence (5’-3’) | Note |
|---|---|---|---|
| **Plasmids construction** | | | |
| PUF_8R/10R (AI-REWIRE) | FLAG-PUF-Nhe1-F | GGCTAGCATGGACTACAAGGACGACGATGACAAGGGTTCT. | this paper |
| | PUF-EcoR1-R | GAATTCCCCTAAGTCAACACCGTTCTTC | |
| hADAR1 | ADAR1-EcoR1-F | GAATTCAAGGCAGAACGCATGGGT | this paper |
| | ADAR1-Not1-R | GCGGCCGCTATACTGGGCAGAGATA | |
| hADAR1-E1008Q | ADAR1-E1008Q-F | CGGACAAGGCACAATCCCTG | this paper |
| | ADAR1-E1008Q-R | CAGGGATTGTGCCTTGTCCG | |
| hADAR2 | hADAR2-EcoR1-F | GAATTCGATCAGACGCCATCTCGCCAG | this paper |
| | hADAR2-Not1-R | GCGGCCGCTTAGGGCGTGAGTGAGAAC | |
| hADAR2-E488Q | ADAR2-E488Q-F | GTCTGGTCAGGGGACGATTCCAG | this paper |
| | ADAR2-E488Q-R | CCAGACTCTATTTTGGTCCGTAGC | |
| PUF_8R/10R (CU-REWIRE) | XTEN-PUF-F | CCGCCACACCCGAAAGTGGCAGGAGCAGGCTTTTG | this paper |
| | T3 promoter-R | ATTAACCCTCACTAAAGG | |
| APOBEC3A | APOBEC3A-Assembly-F | TAATACGACTCACTATAGGGAGAGCCGCCACCATGGAAGC( | this paper |
| | APOBEC3A-XTEN-R | CAAAAGCCTGCTCCTGCCACTTTCGGGTGTGGCGG | |
| APOBEC3A-E72A | APOBEC3A-E72A-F | GCGGCTCTGCGCTTCTTGG | this paper |
| | APOBEC3A-E72A-R | GGTCCAAGAAGCGCAGAGCC | |
| EGFP | EGFP-BamH1-F | CGGGATCCACCATGGTGAGCAAGGGCGAG | this paper |
| | EGFP-Not1-R | ATTTGCGGCCGCTCACTTGTCATCGTCG | |
| EGFP K163X | EGFP K163X-mut-F | AAGCAGAAGAACGGCATCTAGGTG | this paper |
| | EGFP K163X-mut-R | GATGCCGTTCTTCTGCTTGTC | |
| EGFP U503C | EGFP-U503C-mut-F | GAACTTCAAGACCCGCCAC | this paper |
| | EGFP-U503C-mut-R | GATGTTGTGGCGGGTCTTG | |
| EGFP U503A | EGFP-U503A-mut-F | GAACTTCAAGAACCGCCAC | this paper |
| | EGFP-U503A-mut-R | GATGTTGTGGCGGTTCTTG | |
| EGFP U503G | EGFP-U503G-mut-F | GAACTTCAAGAGCCGCCAC | this paper |
| | EGFP-U503G-mut-R | GATGTTGTGGCGGCTCTTG | |
| GRIA2-minigene | GRIA2-Nhe1-F | GCTAGCAAAGCTGATATTGCAATTG | this paper |
| | GRTA2-Not1-R | CGCGGCCGCTTACCTGAAAAACTCTTTAGTG | |
| **AAV9 vector construction** | | | |
| pAV-AI-REWIRE 4.1 | FLAG+PUF-Nhe1-F | GGCTAGCATGGACTACAAGGACGACGATGACAAGGGTTCT. | this paper |
| | AAV9-ADAR2-R | AGATCCGGTGGATCGGATATCTTAGGGCGTGAGTGAGAA CTG | |
| pAV-CU-REWIRE 3.0 | T7 promoter-F | TAATACGACTCACTATAGGG | this paper |
| | AAV9-Flag-R | GATCCGGTGGATCGGATATCTTACTTGTCATCGTCGTCCT TG | |
| pAV-PUF Control | pAV-PUF-Assembly-F | CTCACGCTTAACTAGCTAGCATGGACTACAAGGACGACGA TGACAAG | this paper |
| | pAV-PUF-Assembly-R | TAGTAGCTCCGCTTCCACGCGTCGGTCCGAATTCCCCTAA GTCAACACCG | |
| **Sanger-seq** | | | |
| EGFP | EGFP-seq-F | ATGGTGAGCAAGGGCGAGGAG | this paper |
| | EGFP-seq-R | CTTGTACAGCTCGTCCATGCC | |
| GRIA2 | GRIA2-seq-F | AAAGCTGATATTGCAATTG | this paper |
| | GRIA2-seq-R | CCTGAAAAACTCTTTAGTG | |
| TP53 | TP53-seq-F | TTCCGGGTCACTGCCATGGAG | this paper |
| | TP53-seq-R | GAACAAGAAGTGGAGAATGTCAG | |
| PPIB | PPIB--seq-F | ATGCTGCGCCTCTCCGAACG | this paper |
| | PPIB-seq-R | TACTCCTTGGCGATGGCAAAGG | |
| BMPR2 | BMPR2--seq-F | CCGAACCCTCTCTTGATCTAG | this paper |
| | BMPR2-seq-R | TGTCTACTTGTTTCAAAGCTG | |
| CTNNB1 | CTNNBI-seq-F | GAACTGTCTTTGGACTCTCAGG | this paper |
| | CTNNBI-seq-R | TTCGGTTGTGAACATCCCGAG | |
| EZH2 | PCDH-minigene-F | gaagattctagcatgGCTAGCATG | this paper |
| | PCDH-minigene-R | tcgcagatccttcgcggccgc | |
| SCN1A | PCDH-minigene-F | gaagattctagcatgGCTAGCATG | this paper |

| | | | | |
|---|---|---|---|---|
| | | PCDH-minigene-R | tcgcagatccttcgcggccgc | |
| | Membrane protein (SARA-Cov-2) | M protein seq-F | ATGGCAGATTCCAACGGTAC | this paper |
| | | M protein seq-R | CTGTACAAGCAAAGCAATATTG | |
| | **Deep-seq** | | barcode + primes | |
| | GRIA2 | GRIA2-Seq-F1 | GCTAGCgagaaacacaaagtagtg | for deep sequencing |
| | | GRIA2-Seq-R1 | ctacagtcaggaaggcagCGATCG | |
| | | GRIA2-Seq-F2 | GAATTCgagaaacacaaagtagtg | |
| | | GRIA2-Seq-R2 | ctacagtcaggaaggcagCTTAAG | |
| Primers | | GRIA2-Seq-F3 | GGATCCgagaaacacaaagtagtg | |
| | | GRIA2-Seq-R3 | ctacagtcaggaaggcagCCTAGG | |
| | | GRIA2-Seq-F4 | TCTAGAgagaaacacaaagtagtg | |
| | | GRIA2-Seq-R4 | ctacagtcaggaaggcagAGATCT | |
| | | GRIA2-Seq-F5 | AAGCTTgagaaacacaaagtagtg | |
| | | GRIA2-Seq-R5 | ctacagtcaggaaggcagTTCGAA | |
| | | GRIA2-Seq-F6 | ACGCGTgagaaacacaaagtagtg | |
| | | GRIA2-Seq-R6 | ctacagtcaggaaggcagTGCGCA | |
| | | GRIA2-Seq-F7 | ACTAGTgagaaacacaaagtagtg | |
| | | GRIA2-Seq-R7 | CTACAGTCAGGAAGGCAGTGATCA | |
| | EGFP | GFP-Seq-F1 | GCTAGCGAACCGCATCGAGCTGAAG | for deep sequencing |
| | | GFP-Seq-F2 | GAATTCGAACCGCATCGAGCTGAAG | |
| | | GFP-Seq-F3 | GGATCCGAACCGCATCGAGCTGAAG | |
| | | GFP-Seq-F4 | TCTAGAGAACCGCATCGAGCTGAAG | |
| | | GFP-Seq-F5 | AAGCTTGAACCGCATCGAGCTGAAG | |
| | | GFP-Seq-F6 | ACGCGTGAACCGCATCGAGCTGAAG | |
| | | GFP-Seq-R1 | GGGTGTTCTGCTGGTAGTGCGATCG | |
| | | GFP-Seq-R2 | GGGTGTTCTGCTGGTAGTGCTTAAG | |
| | | GFP-Seq-R3 | GGGTGTTCTGCTGGTAGTGCCTAGG | |
| | | GFP-Seq-R4 | GGGTGTTCTGCTGGTAGTGAGATCT | |
| | | GFP-Seq-R5 | GGGTGTTCTGCTGGTAGTGTTCGAA | |
| | | GFP-Seq-R6 | GGGTGTTCTGCTGGTAGTGTGCGCA | |
| | TP53 | TP53-Seq-F1 | GCTAGCATGGCCATCTACAAGCAG | for deep sequencing |
| | | TP53-Seq-F2 | GAATTCATGGCCATCTACAAGCAG | |
| | | TP53-Seq-F3 | GGATCCATGGCCATCTACAAGCAG | |
| | | TP53-Seq-R1 | GTCAGAGCCAACCTCAGGCGATCG | |
| | | TP53-Seq-R2 | GTCAGAGCCAACCTCAGGCTTAAG | |
| | | TP53-Seq-R3 | GTCAGAGCCAACCTCAGGCCTAGG | |
| | PPIB | PPIB-Seq-F1 | GCTAGCTGGCCTTAGCTACAGGAG | for deep sequencing |
| | | PPIB-Seq-F2 | GAATTCTGGCCTTAGCTACAGGAG | |
| | | PPIB-Seq-F3 | GGATCCTGGCCTTAGCTACAGGAG | |
| | | PPIB-Seq-R1 | TGCGTTGGCCATGCTCACCGATCG | |
| | | PPIB-Seq-R2 | TGCGTTGGCCATGCTCACCTTAAG | |
| | | PPIB-Seq-R3 | TGCGTTGGCCATGCTCACCCTAGG | |
| | DMD | DMD-Seq-F1 | GCTAGCGTGGAGATCACGCAACTG | for deep sequencing |
| | | DMD-Seq-F2 | GAATTCGTGGAGATCACGCAACTG | |
| | | DMD-Seq-F3 | GGATCCGTGGAGATCACGCAACTG | |
| | | DMD-Seq-R1 | TAGCATCTTCTTTTTCTGCGATCG | |
| | | DMD-Seq-R2 | TAGCATCTTCTTTTTCTGCTTAAG | |
| | | DMD-Seq-R3 | TAGCATCTTCTTTTTCTGCCTAGG | |
| | COL3A1 | COL3A1-Seq-F1 | GCTAGCCTCCATAGAAGATTctaG | for deep sequencing |
| | | COL3A1-Seq-F2 | GAATTCCTCCATAGAAGATTctaG | |
| | | COL3A1-Seq-F3 | GGATCCCTCCATAGAAGATTctaG | |
| | | COL3A1-Seq-R1 | CAGATGGACCTATAGCACCGATCG | |
| | | COL3A1-Seq-R2 | CAGATGGACCTATAGCACCTTAAG | |
| | | COL3A1-Seq-R3 | CAGATGGACCTATAGCACCCTAGG | |
| | HBB | HBB-Seq-F | CATGGCTAGCATGGTGCATC | for deep sequencing |

| | | | | |
|---|---|---|---|---|
| | | HBB-Seq-R | CATGAGCCTTCACCTTAGGG | |
| | EZH2 | EZH2-Seq-F1 | GCTAGCATGAAGGGTAACAAAATTCG | for deep sequencing |
| | | EZH2-Seq-F2 | GAATTCATGAAGGGTAACAAAATTCG | |
| | | EZH2-Seq-F3 | GGATCCATGAAGGGTAACAAAATTCG | |
| | | EZH2-Seq-R1 | gcTTAAGGGATTTCCATTTCCGATCG | |
| | | EZH2-Seq-R2 | gcTTAAGGGATTTCCATTTCCTTAAG | |
| | | EZH2-Seq-R3 | gcTTAAGGGATTTCCATTTCCCTAGG | |
| | SCN1A | SCN1A-Seq-F1 | GCTAGCGAAGGCTGGAATATCTTTG | for deep sequencing |
| | | SCN1A-Seq-F2 | GAATTCGAAGGCTGGAATATCTTTG | |
| | | SCN1A-Seq-F3 | GGATCCGAAGGCTGGAATATCTTTG | |
| | | SCN1A-Seq-R1 | GACGAGGGTTAAATTTCCCGATCG | |
| | | SCN1A-Seq-R2 | GACGAGGGTTAAATTTCCCTTAAG | |
| | | SCN1A-Seq-R3 | GACGAGGGTTAAATTTCCCCTAGG | |
| | SCN5A | SCN5A-Seq-F1 | GCTAGCGCATGCCTTCCTCATCATC | for deep sequencing |
| | | SCN5A-Seq-F2 | GAATTCGCATGCCTTCCTCATCATC | |
| | | SCN5A-Seq-F3 | GGATCCGCATGCCTTCCTCATCATC | |
| | | SCN5A-Seq-R1 | CTCATCAGGGGCTGTGAGCGATCG | |
| | | SCN5A-Seq-R2 | CTCATCAGGGGCTGTGAGCTTAAG | |
| | | SCN5A-Seq-R3 | CTCATCAGGGGCTGTGAGCCTAGG | |

### Minigenes

| | | | | |
|---|---|---|---|---|
| | OTC singal | | ATGTTGTTCAACTTGCGCATCCTGTTGAACAATGCAGCCTT TAGGAACGGTCACAACTTCATGGTTCGCAACTTCCGGTGT GGGCAGCCACTGCAAAACAAAGTGCAG | this paper |
| | T2A-mcherry-minigene | | GTCGACGGAAGCGGAGAGGGCAGAGGAAGTCTGCTAACA TGCGGTGACGTCGAGGAGAATCCTGGACCTATGGTAAGTA AGGGCGAGGAGG | this paper |
| | Minigene-PUF-NLS-FLAG | | GTACTACATGAAGAACGGTGTTGACTTAGGGGCTAGCGGA TCCCCCAAGAAGAAGAGGAAAGTCTCTAGAGACTACAAGG ACGACGATGACAAGTAAGCGGCCGCTTCCCTTTAGTGAGG GTTAA | this paper |
| | NM_004006.2(DMD):c.1652G>A (p.Trp551Ter) | | CCATAGAAGATTCTAGCATGGCTAGCGTGGTAGTTGATGA ATCTAGTGGAGATCACGCAACTGCTGCTTTGGAAGAACAA CTTAAGGTATTGGGAGATCGATGGGCAAACATCTGTAGAT GGACAGAAGACCGCTAGGTTCTTTTACAAGACATCCTTCT CAAATGGCAACGTCTTACTGAAGAACAGTGCCTTTTTAGTG CATGGCTTTCAGAAAAAGAAGATGCTAAGCGAAGGATCTG CGATCGCTC | this paper |
| | NM_004006.2(DMD):c.1682G>A (p.Trp561Ter) | | CCATAGAAGATTCTAGCATGGCTAGCGTGGTAGTTGATGA ATCTAGTGGAGATCACGCAACTGCTGCTTTGGAAGAACAA CTTAAGGTATTGGGAGATCGATGGGCAAACATCTGTAGAT GGACAGAAGACCGCTGGGTTCTTTTACAAGACATCCTTCT CAAATAGCAACGTCTTACTGAAGAACAGTGCCTTTTTAGTG CATGGCTTTCAGAAAAAGAAGATGCTAAGCGAAGGATCTG CGATCGCTC | this paper |
| | NM_000518.4(HBB):c.47G>A (p.Trp16Ter) | | AACCTCAAACAGACACCATGGTGCATCTGACTCCTGAGGA GAAATCTGCCGTTACTGCCCTGTAGGGCAAGGTGAACGTG GATGAAGTTGGTGGTGAGGCCCTGGGCAGGCTGCTGGTG GTCTACCCTTGGACCCAGAGGTTCTTTGAGTCCTTTGGGG ATCTGTCCACTCCTGATGCTGTTATGGGCAACCCTAAGGT GAAGGCTCATGGCAAGAAAGGCTCGCTTTCTTGCT | this paper |
| | NM_000518.4(HBB):c.113G>A (p.Trp38Ter) | | AACCTCAAACAGACACCATGGTGCATCTGACTCCTGAGGA GAAATCTGCCGTTACTGCCCTGTGGGGCAAGGTGAACGT GGATGAAGTTGGTGGTGAGGCCCTGGGCAGGCTGCTGGT GGTCTACCCTTAGACCCAGAGGTTCTTTGAGTCCTTTGGG GATCTGTCCACTCCTGATGCTGTTATGGGCAACCCTAAGG TGAAGGCTCATGGCAAGAAAGGCTCGCTTTCTTGCT | this paper |
| Minigenes | NM_000090.3(COL3A1):c.565G>A (p.Gly189Ser) | | CTCCATAGAAGATTCTAGAATGGGCCCTCCCGGTCCCCCT GGTACATCTAGTCATCCTGGTTCCCCTGGATCTCCAGGAT ACCAAGGACCCCCTGGTGAACCTGGGCAAGCTGGTCCTT CAGGCCCTCCAGGACCTCCTGGTGCTATAGGTCCATCTG GTCCTGCTGGAAAAGATGGAGAATCAGGTAGACCCGGAC GACCTGGAGAGCGAGGATTGCCTGGACCTCCAGGTGCGG CCGCGAAGGATCTG | this paper |

| | | | |
|---|---|---|---|
| NM_000090.3(COL3A1):c.637G>A (p.Gly213Ser) | | CTCCATAGAAGATTCTAGAATGGGCCCTCCCGGTCCCCCT<br>GGTACATCTGGTCATCCTGGTTCCCCTGGATCTCCAGGAT<br>ACCAAGGACCCCCTGGTGAACCTGGGCAAGCTGGTCCTT<br>CAAGCCCTCCAGGACCTCCTGGTGCTATAGGTCCATCTGG<br>TCCTGCTGGAAAAGATGGAGAATCAGGTAGACCCGGACG<br>ACCTGGAGAGCGAGGATTGCCTGGACCTCCAGGTGCGGC<br>CGCGAAGGATCTG | this paper |
| NM_004456.4(EZH2):c.2191T>C (p.Tyr731His) | | GAAGATTCTAGCATGGCTAGCATGAAGGGTAACAAAATTC<br>GTTTTGCAAATCATTCGGTAAATCCAAACTGCTATGCAAAA<br>GTTATGATGGTTAACGGTGATCACAGGATAGGTATTTTTGC<br>CAAGAGAGCCATCCAGACTGGCGAAGAGCTGTTTTTTGAT<br>CACAGATACAGCCAGGCTGATGCCCTGAAGTATGTCGGCA<br>TCGAAAGAGAAATGGAAATCCCTTAAGCGGCCGCGAAGGA<br>TCTGCGAT | this paper |
| NM_001165963.1(SCN1A):c.2588T>C (p.Ser863Leu) | | GAAGATTCTAGCATGGCTAGCATGGAAGGCTGGAATATCT<br>TTGACGGTTTTATTGTGACGCTTAGCCTGGTAGAACTTGG<br>ACTCGCCAATGTGGAAGGATTATCTGTTCTCCGTTCATTTC<br>GATCGCTGCGAGTTTTCAAGTTGGCAAAATCTTGGCCAAC<br>GTTAAATATGCTAATAAAGATCATCGGCAATTCCGTGGGG<br>GCTCTGGGAAATTTAACCCTCGTCTAAGCGGCCGCGAAG<br>GATCTGCGAT | this paper |
| NM_198056.2(SCN5A):c.2756T>C (p.Phe919Ser) | | GAAGATTCTAGCATGGCTAGCATGCATGCCTTCCTCATCA<br>TCTTCCGCATCCTCTGTGGAGAGTGGATCGAGACCATGTG<br>GGACTGCATGGAGGTGTCGGGGCAGTCATTATGCCTGCT<br>GGTCTCCTTGCTTGTTATGGTCATTGGCAACCTTGTGGTC<br>CTGAATCTCTTCCTGGCCTTGCTGCTCAGCTCCTTCAGTG<br>CAGACAACCTCACAGCCCCTGATGAGTAAGCGGCCGCGA<br>AGGATCTGCGAT | this paper |
| Membrane protein (SARA-Cov-2) | 2019-nCoV-5 in pUC57-Amp | ATGGCAGATTCCAACGGTACTATTACCGTTGAAGAGCTTA<br>AAAAGCTCCTTGAACAATGGAACCTAGTAATAGGTTTCCTA<br>TTCCTTACATGGATTTGTCTTCTACAATTTGCCTATGCCAA<br>CAGGAATAGGTTTTTGTATATAATTAAGTTAATTTTCCTCTG<br>GCTGTTATGGCCAGTAACTTTAGCTTGTTTTGTGCTTGCTG<br>CTGTTTACAGAATAAATTGGATCACCGGTGGAATTGCTATC<br>GCAATGGCTTGTCTTGTAGGCTTGATGTGGCTCAGCTACT<br>TCATTGCTTCTTTCAGACTGTTTGCGCGTACGCGTTCCAT<br>GTGGTCATTCAATCCAGAAACTAACATTCTTCTCAACGTGC<br>CACTCCATGGCACTATTCTGACCAGACCGCTTCTAGAAAG<br>TGAACTCGTAATCGGAGCTGTGATCCTTCGTGGACATCTT<br>CGTATTGCTGGACACCATCTAGGACGCTGTGACATCAAGG<br>ACCTGCCTAAAGAAATCACTGTTGCTACATCACGAACGCTT<br>TCTTATTACAAATTGGGAGCTTCGCAGCGTGTAGCAGGTG<br>ACTCAGGTTTTGCTGCATACAGTCGCTACAGGATTGGCAA<br>CTATAAATTAAACACAGACCATTCCAGTAGCAGTGACAATA<br>TTGCTTTGCTTGTACAG | GENEWIZ (HA3620-5/T504644) |

### Plasmids used in this study

| Reagent/Resource | source |
|---|---|
| pGL-PUF | this paper |
| pGL-PUF-10R | this paper |
| pGL-ADAR1 | this paper |
| pGL-ADAR2 | this paper |
| pGL-PUF-ADAR1 | this paper |
| pGL-PUF-ADAR1-E1008Q | this paper |
| pGL-PUF-ADAR2 | this paper |
| pGL-PUF-ADAR2-E488Q | this paper |
| pGL-PUF10R-ADAR2 | this paper |
| pGL-PUF10R-ADAR2-E488Q | this paper |
| pGL-PUF10R*-ADAR2 | this paper |
| pGL-PUF10R*-ADAR2-E488Q | this paper |
| pGL-A3A | this paper |
| pGL-A3A-PUF-NLS | this paper |
| pGL-A3A-PUF | this paper |
| pGL-A3A-E72A-PUF | this paper |
| pGL-A3A-PUF10R | this paper |
| pGL-A3A-PUF10R* | this paper |
| pCDH-EGFP | this paper |
| pCDH-GRIA2 | this paper |
| pCDH-DMD G1652A | this paper |
| pCDH-DMD G1682A | this paper |

| | |
|---|---|
| pCDH-COL3A1 G565A | this paper |
| pCDH-COL3A1 G637A | this paper |
| pCDH-HBB G47A | this paper |
| pCDH-HBB G113A | this paper |
| pCDH-EZH2 T2191C | this paper |
| pCDH-SCN1A T2588C | this paper |
| pCDH-SCN5A T2756C | this paper |

**REWIRE plasmids used in this study.**

| Figure | Target gene | REWIREs | Recognition site | source |
|---|---|---|---|---|
| 1A, S1C-F | EGFP | AI-REWIRE (EGFP-A437) | CGUCUAUA | this paper |
| 1F, S3A | GRIA2 | AI-REWIRE (GRIA2-A1820)-6 | GAUGCGAU | this paper |
| 1F, S3A | GRIA2 | AI-REWIRE (GRIA2-A1820)-7 | AUGCGAUA | this paper |
| 1F, S3A | GRIA2 | AI-REWIRE (GRIA2-A1820)-8 | UGCGAUAU | this paper |
| 1F, S3A | GRIA2 | AI-REWIRE (GRIA2-A1820)-9 | GCGAUAUU | this paper |
| 1F, S3A | GRIA2 | AI-REWIRE (GRIA2-A1820)-10 | CGAUAUUU | this paper |
| 1F, S3A | GRIA2 | AI-REWIRE (GRIA2-A1820)-11 | GAUAUUUC | this paper |
| 1F, S3A | GRIA2 | AI-REWIRE (GRIA2-A1820)-12 | AUAUUUCG | this paper |
| 1F, S3A | GRIA2 | AI-REWIRE (GRIA2-A1820)-13 | UAUUUCGC | this paper |
| 2A-F, S4A-B | EGFP K163X | AI-REWIRE (EGFP-A488) | CUUCAAGA | this paper |
| 2G-H | TP53 | AI-REWIRE (TP53-A553) | UCUGGCCC | this paper |
| 2J | TP53 | AI-REWIRE (TP53-A512) | GGCGCUGC | this paper |
| 2J,S4C | PPIB | AI-REWIRE (PPIB-A325) | UGGCACAG | this paper |
| 2J | PPIB | AI-REWIRE (PPIB-A123) | UCAAGGUG | this paper |
| 2J | BMPR2 | AI-REWIRE (BMPR2-A898) | GCCGUCUU | this paper |
| 2J | DMD | AI-REWIRE (DMD-1682) | UUACUGAA | this paper |
| 2J,S4E | DMD | AI-REWIRE (DMD-1652) | UUACAAGA | this paper |
| 2J | COL3A1 | AI-REWIRE (COL3A1-565) | UGGUUCCC | this paper |
| 2J | HBB | AI-REWIRE (HBB-47) | UGAACGUG | this paper |
| 2J | HBB | AI-REWIRE (HBB-113) | GUUCUUUG | this paper |
| 3B, 3F | EGFP | CU-REWIRE (EGFP-C504) | CUUCAAGA | this paper |
| 3D | EGFP | CU-REWIRE (EGFP)-4 | AACUUCAA | this paper |
| 3D | EGFP | CU-REWIRE (EGFP)-3 | ACUUCAAG | this paper |
| 3D | EGFP | CU-REWIRE (EGFP)-1 | UUCAAGAU | this paper |
| 3D | EGFP | CU-REWIRE (EGFP)-0 | UCAAGAUC | this paper |
| 3G-H | CTNNB1 | CU-REWIR (CTNNB1-C1549) | AUUCGAAA | this paper |
| S5B | EZH2 | CU-REWIR (EZH2-C2191) | UUUUUUGA | this paper |
| S5C | SCN1A | CU-REWIR (SCN1A-C2588) | CAUUUCGA | this paper |
| S5D | M (SARS-Cov2) | CU-REWIR (M-C246) | AAUUGCUA | this paper |
| 4B, 4H | EGFP | AI-REWIRE 3/4 (EGFP-A437) | CGUCUAUAUC | this paper |
| 4E, 4H | EGFP | CU-REWIRE 2/3(EGFP-C459) | AACGUCUAUA | this paper |
| S6C | PPIB | AI-REWIRE 4 (PPIB-A325) | AUGGCACAGG | this paper |
| S6F | EZH2 | CU-REWIR (EZH2-C2191) | UGUUUUUUGA | this paper |
| S6G | SCN1A | CU-REWIR (SCN1A-C2588) | UUCAUUUCGA | this paper |
| 5B | EGFP | AI-REWIRE4.1 (EGFP-A437) | CGUCUAUAUC | this paper |
| 5D | EGFP | CU-REWIRE3.0 (EGFP-C459) | AACGUCUAUA | this paper |
