## Supplementary material for "Programmable RNA base editing with a single gRNA-free enzyme": Table S2

### Table S2 Summary of RNA-seq

| Sample name | Sample Code | library_selection | Mapped bases (Gbp)/reads |
|---|---|---|---|
| HEK 293T -Mock | 19R475 | Inverse rRNA | 43,524,508 |
| | 19R476 | Inverse rRNA | 44,641,176 |
| | 19R170 | Inverse rRNA | 66,163,132 |
| PUF-Ctrl | 19R406 | Inverse rRNA | 43,374,442 |
| | 19R407 | Inverse rRNA | 44,819,178 |
| | 19R412 | Inverse rRNA | 46,625,470 |
| ADAR1-Ctrl | 18R347 | Inverse rRNA | 65,880,175 |
| ADAR2-Ctrl | 18R348 | Inverse rRNA | 53,197,578 |
| AI-REWIRE1.0 | 19R542 | Inverse rRNA | 59,863,876 |
| | 19R544 | Inverse rRNA | 63,341,064 |
| | 19R546 | Inverse rRNA | 58,452,259 |
| AI-REWIRE1.1 | 19R543 | Inverse rRNA | 57,618,253 |
| | 19R545 | Inverse rRNA | 60,230,139 |
| | 19R547 | Inverse rRNA | 60,700,971 |
| AI-REWIRE2.0 | 19R166 | Inverse rRNA | 59,649,571 |
| | 19R410 | Inverse rRNA | 46,734,698 |
| | 19R404 | Inverse rRNA | 45,024,076 |
| AI-REWIRE2.1 | 19R405 | Inverse rRNA | 43,653,730 |
| | 19R411 | Inverse rRNA | 48,636,350 |
| | 19R167 | Inverse rRNA | 71,387,707 |
| CU-REWIRE-fig3 | 19R548 | Inverse rRNA | 62,645,030 |
| | 19R550 | Inverse rRNA | 61,471,374 |
| | 19R200 | Inverse rRNA | 54,116,119 |
| Mut.CU-REWIRE | 19R552 | Inverse rRNA | 57,709,577 |
| | 19R553 | Inverse rRNA | 56,506,040 |
| | 19R554 | Inverse rRNA | 65,357,307 |
| PUF-10R-Ctrl | 19R713 | Inverse rRNA | 55,319,830 |
| | 19R714 | Inverse rRNA | 61,045,956 |
| | 19R715 | Inverse rRNA | 67,160,143 |
| AI-REWIRE3.0 | 19R402 | Inverse rRNA | 45,625,237 |
| | 19R164 | Inverse rRNA | 37,679,509+21834460 |
| | 19R408 | Inverse rRNA | 46,386,481 |
| AI-REWIRE3.1 | 19R403 | Inverse rRNA | 48,010,909 |
| | 19R165 | Inverse rRNA | 39,801,091+8833524 |
| | 19R409 | Inverse rRNA | 44,491,764 |
| AI-REWIRE4.1 | 20R354 | Inverse rRNA | 59,434,200 |
| | 20R355 | Inverse rRNA | 58,012,481 |
| | 20R356 | Inverse rRNA | 54,261,250 |
| CU-REWIRE1.0 | 19R847 | Inverse rRNA | 52,220,507 |
| | 19R848 | Inverse rRNA | 54,581,004 |
| | 19R849 | Inverse rRNA | 55,577,206 |
| CU-REWIRE2.0 | 19R850 | Inverse rRNA | 50,633,163 |
| | 19R851 | Inverse rRNA | 59,300,854 |
| | 19R852 | Inverse rRNA | 52,490,588 |
| CU-REWIRE3.0 | 20R317 | Inverse rRNA | 40,228,258+16213949 |
| | 20R318 | Inverse rRNA | 34,608,441+19669145 |
| | 20R319 | Inverse rRNA | 37,243,589+15402573 |
| PUF-Ctrl-mouse | 20R576 | Inverse rRNA | 56,037,126 |
| | 20R577 | Inverse rRNA | 48,361,326 |
| | 20R578 | Inverse rRNA | 56,335,386 |
| AI-REWIRE4.1-mouse | 20R570 | Inverse rRNA | 50,969,411 |
| | 20R571 | Inverse rRNA | 45,647,452 |
| | 20R572 | Inverse rRNA | 47,556,677 |
| CU-REWIRE3.0-mouse | 20R573 | Inverse rRNA | 64,660,319 |
| | 20R574 | Inverse rRNA | 51,067,782 |
| | 20R575 | Inverse rRNA | 50,168,937 |
| AI-REWIRE2.0-PPIB | 20R304 | Inverse rRNA | 54,347,576 |
| | 20R305 | Inverse rRNA | 59,308,701 |
| | 20R306 | Inverse rRNA | 62,697,457 |
| AI-REWIRE4.0-PPIB | 20R307 | Inverse rRNA | 66,110,282 |
| | 20R308 | Inverse rRNA | 58,890,188 |
| | 20R293 | Inverse rRNA | 56,633,186 |
